# Dose-additive and dose-dependent fetal testis toxicity of a nine-phthalate mixture in the rat

**DOI:** 10.64898/2026.10.11.755118

**Authors:** Joceline Helmbreck, Ziqi Zhao, Poonam Mehta, Patrick Schupp, Aditi Marehalli, Jason Scott, Matthew J. Woodcock, Andre Gomes, Cindy Su, Matthew Manwaring, Claire Gould, Thais Del Rosario Hernandez, Petra den Hollander, Susan J. Hall, Adina Badea, Sendurai Mani, Kurt D. Pennell, Daniel J. Spade

## Abstract

Phthalates (ortho-phthalic acid diesters) are used in a variety of manufacturing processes, particularly as non-covalent plasticizers in flexible polyvinyl chloride. Phthalates are male reproductive toxicants that disrupt the development of the fetal testis, resulting in malformed seminiferous cords, multinucleated germ cells (MNGs), and decreased testosterone production. We previously reported that MNGs were a sensitive, dose-dependent marker of dipentyl phthalate (DPEP) germ cell toxicity. To test whether germ cell effects of phthalates are dose-additive, we measured the induction of MNGs by a phthalate mixture. We treated timed pregnant Sprague Dawley rats with an equipotent mixture of nine phthalates, spanning a two-log dose range, from GD 17-21. We found that the phthalate mixture effects on morphogenesis of the testis, assessed through histological analysis of MNGs and seminiferous cord diameter were dose-additive. Consistent with prior results, we also found that the testosterone-reducing effect of the phthalate mixture was dose-additive and dose-dependent. Induction of MNGs had the lowest effective dose of any endpoint that we measured, with a benchmark dose (BMD_10_) of 9.402 (6.093, 13.13) mg (∑ all phthalates)/kg/d, which is equivalent to a BMD_10_ of 3.517 (2.275, 4.919) mg DPEP/kg/d, based on relative potency factors. Quantitative analysis of phthalate metabolites in dam serum showed that the metabolites were present in approximately the same relative concentrations as the parent compounds in the dosing mixture, but with lower levels of monocyclohexyl phthalate and monobenzyl phthalate. Proteomic and spatial transcriptomic analyses showed significant decreases in expression of Leydig cell genes and proteins involved in steroid hormone biosynthesis, as well as enrichment of proteins related to metabolism of lipids and amino acids. Proteomic BMD analysis produced a lowest BMDmedian of 2.156 (0.204, 27.830) mg (∑ all phthalates)/kg/d, which was within a 4.36-fold margin of the lowest apical endpoint BMD.

## Introduction

Ortho-phthalic acid diesters (phthalates) are a class of chemicals used to plasticize polyvinyl chloride and also used in a variety of industrial products, manufactured goods, and personal care products ^1, 2^. Humans are exposed to low doses of phthalates from a variety of sources, with the highest exposure to certain phthalates in medical settings when patients receive blood transfusions and total parenteral nutrition ^3-6^. Phthalates are well-characterized male reproductive toxicants ^7^. The risk of phthalate toxicity, particularly to the developing male reproductive tract ^8^, has led the U.S. EPA to initiate a cumulative risk assessment process for phthalates ^9^.

Phthalates disrupt the development of the fetal testis, resulting in several well-characterized effects that can be categorized as anti-androgenic effects and seminiferous cord effects ^7^. The anti-angiogenic effects of phthalates result from a decrease in fetal testicular testosterone production ^10^, which occurs as early as 1-6 hours after phthalate exposure ^11^ and leads to decreased masculinization of the male reproductive tract ^12, 13^. These anti-androgenic effects are caused by phthalates with parent aliphatic side chains between 3 and 8 carbons in length ^10^. Phthalates also alter the morphogenesis of the fetal testis, adversely affecting seminiferous cord development and the development of fetal Sertoli cells and germ cells. These effects, sometimes referred to as “testicular dysgenesis” ^14^, can include reduced Sertoli cell mitosis and altered Sertoli cell cytoskeleton ^15, 16^, ruptured cords and ectopic cells ^14^, abnormally large cord diameter, and induction of multinucleated germ cells (MNGs) ^17, 18^. Like the reduction in testosterone, MNGs are also induced by phthalates with medium-length aliphatic side chains ^19^. However, MNGs can be induced without a significant decrease in testosterone, and reduced testosterone production is not sufficient to induce MNGs ^20-22^. It is likely that there is a single, as-yet unknown, molecular initiating event for toxic effects of phthalates on both the production of androgens and the morphogenesis of the seminiferous cords.

Because the seminiferous cord effects of phthalates are relevant to humans and can occur independently of changes in testosterone production ^20^, we have sought to characterize dose-response for phthalate-induced MNGs, which are a quantitative metric of phthalate-induced seminiferous cord effects. We recently reported that dipentyl phthalate (DPeP) induced MNGs in the fetal rat testis with a benchmark dose (BMD_10_) of 2.675 mg/kg/d, which was the lowest BMD_10_ for any endpoint that we tested ^23^, including testosterone production. It has previously been demonstrated that the anti-androgenic effects of phthalates are dose-additive ^24-28^. Here, to test whether the effects of phthalates on the seminiferous cord are also dose additive, we exposed timed pregnant Sprague Dawley rats to a mixture of 9 phthalates from GD 17-21, using a mixture that was designed to be equipotent for the better-characterized anti-androgenic effects ^10, 24^. We compare the effects of the mixture on germ cell and seminiferous cord development with effects on testosterone production, and we test whether these effects can be modeled by the dose-addition equation.

## Methods

### Animals

All procedures involving animals were approved by the Brown University Institutional Animal Care and Use Committee and were performed in accordance with the Guide for the Care and Use of Laboratory Animals. Euthanasia methods are approved by the American Veterinary Medical Association. Timed pregnant Sprague Dawley rats (Crl:CD(SD), strain code 001) were obtained from Charles River Laboratories (Shrewsbury, MA) and arrived at Brown University on gestation day (GD) 12. The day on which a copulatory plug was detected was considered to be GD 1. Rats were housed in polysulfone Optirat cages (Animal Care Systems, Centennial, CO, USA) with autoclaved hardwood bedding (Sani-Chips) and had free access to LabDiet 5010 rodent diet and RO water, chlorinated at 1-3 ppm. The animal care facility is on a 12 hr light:12 hr dark cycle with lights on at 7:00 ET and is maintained at 20-22.2°C and 30-70% humidity, with a set point of 50%.

### Chemicals and Experimental Procedures

A mixture of nine phthalates was generated as described in Table 1. Briefly, an equipotent mixture of diisobutyl phthalate (DIBP), di-*n*-butyl phthalate (DBP), benzyl butyl phthalate (BBP), dipentyl phthalate (DPEP), dicyclohexyl phthalate (DCHP), dihexyl phthalate (DHP), diisoheptyl phthalate (DIHEP), di-(2-ethylheyxl) phthalate (DEHP), and diheptyl phthalate (DHP) was mixed in corn oil vehicle (V). Potency was based on ED_50_ for reduction of T production as reported by Furr, Lambright, Wilson, Foster and Gray ^10^. The initial mixture was the highest study dose (A), and each subsequent dose (B, C, D, and E in descending order) was a ½-log dilution, made by serial dilution in corn oil vehicle. Chemicals were mixed by weight using an analytical balance and stirred in stoppered borosilicate glass Erlenmeyer flasks with Teflon-coated magnetic stir bars. To the greatest extent possible, dosing solutions were protected from light exposure by wrapping flasks in aluminum foil while stirring. Dosing solutions were aliquoted for daily use and stored in borosilicate glass scintillation vials, in an opaque cardboard box at room temperature, and were used within one week.

**Table 1.** Chemicals and parameters of 9-phthalate mixture.

| Name of Dose Level |  |  |  |  |  |  |  |  |  | A | B | C | D | E |
| --- | --- | --- | --- | --- | --- | --- | --- | --- | --- | --- | --- | --- | --- | --- |
| phthalate | abbrev | parent C | total C | CASRN | source | PN | lot | purity (%) | ED<sub>50</sub>* (mg/kg/d) | 0.5 log | 1 log | 1.5 log | 2 log | 2.5 log |
| diisobutyl | DIBP | 3 | 4 | 84-69-5 | Sigma Aldrich | 152641 | MKCN5163 | 99.6 | 288.2 | 91.1 | 28.8 | 9.1 | 2.9 | 0.9 |
| di-*n*-butyl | DBP | 4 | 4 | 84-74-2 | Sigma Aldrich | 524980 | MKBL8504V | 99.4 | 157.9 | 49.9 | 15.8 | 5.0 | 1.6 | 0.5 |
| benzyl butyl | BBP | 4, 6 | 4, 6 | 85-68-7 | Sigma Aldrich | 308501 | MKCN9008 | 97.7 | 172.4 | 54.5 | 17.2 | 5.4 | 1.7 | 0.5 |
| dipentyl | DPEP | 5 | 5 | 131-18-0 | TCI America | P0291 | MYZGF-BP | ≥ 98 | 81.6 | 25.8 | 8.2 | 2.6 | 0.8 | 0.3 |
| dicyclohexyl | DCHP | 6 | 6 | 84-61-7 | Sigma Aldrich | 306150 | MKBB1696 | 99.9 | 61.6 | 19.5 | 6.2 | 1.9 | 0.6 | 0.2 |
| dihexyl | DHP | 6 | 6 | 84-75-3 | TCI America | P1344 | ENTQH-CF | 99.6 | 59.2 | 18.7 | 5.9 | 1.9 | 0.6 | 0.2 |
| diisoheptyl | DIHEP | 6 | 7 | 71888-89-6 | TRC | D455720 | 11-KMT-88-1 | 99.34 | 361.6 | 114.3 | 36.2 | 11.4 | 3.6 | 1.1 |
| di-(2-ethylhexyl) | DEHP | 6 | 8 | 117-81-7 | Sigma Aldrich | D201154 | MKCK4506 | 99.7 | 359.8 | 113.7 | 36.0 | 11.4 | 3.6 | 1.1 |
| diheptyl | DHEP | 7 | 7 | 3648-21-3 | Sigma Aldrich | 454818 | BCBZ7192 | 98.4 | 401.7 | 126.9 | 40.2 | 12.7 | 4.0 | 1.3 |
| DPEP equivalent dose (mg/kg)** |  |  |  |  |  |  |  |  |  | 232.1 | 73.5 | 23.2 | 7.3 | 2.3 |
| total phthalate dose (mg/kg) |  |  |  |  |  |  |  |  |  | 614.3 | 194.4 | 61.4 | 19.4 | 6.1 |

Dose levels named A (high)-E (low), calculated as 0.5-2.5 log dilutions from each phthalate ED<sub>50</sub>. All doses listed in mg/kg/d. Lot-specific purity reported from certificate of analysis, except DPEP, where COA was not available. abbrev: abbreviation; parent C: length of parent side chain; total C: total C in side chain, including branches; PN: product number. \*ED<sub>50</sub> for reduction of testosterone production from Furr et al. 2014. \*\*DPeP equivalent dose calculating using ED<sub>50</sub> ratio. Mixed in corn oil vehicle (Sigma Aldrich PN C8267).

Rats were weighed daily to confirm that they were gaining weight and assigned to a treatment group by sorting on a random number assigned by the =RAND() function in Excel. The nine-phthalate mixture or vehicle control was administered by daily oral gavage in a 2 mL/kg solution, using a Hamilton syringe and ball-end gavage needle in the morning. On GD 21, the final dose was administered 1 hour before euthanasia. Rats were euthanized by overdose of inhaled isoflurane in a static isoflurane chamber. Blood was drawn by cardiac puncture, allowed to clot at room temperature, and centrifuged in BD Vacutainer Serum Separation Tubes (red/gray top) with polymer gel (Becton Dickinson, Franklin Lakes, NJ). Fetuses were removed from the uterus and euthanized by decapitation with a clean, disposable razor blade. Fetuses were dissected under a Zeiss Discovery V8 stereomicroscope, and sex was determined by observation of the gonads. Testes were collected from all male fetuses. Testes with attached mesonephroi to be used for histology and immunofluorescence were immediately fixed in modified Davidson’s solution, fixed for 30 minutes at room temperature with agitation, then transferred to 70% EtOH and stored at 4°C. Three testes from different males in each litter were cleanly dissected from the mesonephros and maintained in phosphate buffered saline (PBS) briefly until they could be transferred to assay media for the *ex vivo* T production assay. Testes to be used for proteomics were dissected from the mesonephros and snap-frozen in liquid nitrogen, then stored at -80°C. Testes to be used for spatial transcriptomics were collected with the attached mesonephros and retained on PBS briefly until they could be transferred to Optimal Cutting Temperature (OCT) Compound (Tissue-Tek). OCT blocks were slowly frozen on dry ice and stored at -80°C.

### Histology

After fixation, testes with attached mesonephroi were processed through graded ethanols, followed by three changes of xylene, two changes of hot paraffin wax, then embedded in paraffin wax. They were trimmed to approximately the middle third of the tissue sample, and 5 µm sections were cut and mounted on Fisherbrand Colorfrost Plus microscope slides. For all staining procedures, slides were deparaffinized in three changes of xylene and rehydrated in graded ethanols. For Quantification of MNGs, slides were then stained with hematoxylin and eosin (H&E). Images of H&E-stained slides were obtained at 40x magnification using an Aperio Scanscope CS. MNGs and slide area were quantified by manual annotation by a blinded scorer in ImageScope v 12.2 (Leica Biosystems).

### Immunofluorescence

Tissue sections were deparaffinized and rehydrated as described above and used for immunofluorescent labeling of SOX9 and AMH, according to our previously published methods ^23^. Briefly, following rehydration, slides were subjected to acidic antigen retrieval by immersing in citric acid antigen retrieval solution (Vector Laboratories) in a vegetable steamer for 20 min, then cooled at room temperature for 20 min. Sections were permeabilized using 0.1% Triton X-100 in PBS for 5 min at room temperature. Nonspecific antigens were blocked by applying 5% normal goat serum and 1% bovine serum albumin in PBS for 60 minutes. Primary antibodies for SOX9 (rabbit polyclonal, MilliporeSigma #AB5535, 1:500 dilution) and AMH (mouse monoclonal, Santa Cruz Biotechnology #sc-166752, 1:50 dilution) were diluted in blocking buffer. Slides were incubated in primary antibody solution overnight at 4°C in an opaque, humidified chamber. Secondary antibodies (goat anti-rabbit IgG-AlexaFluor 568 conjugate, ThermoFisher Scientific #A11036, 1:500 dilution; goat anti-mouse IgG-AlexaFluor 488 conjugate, ThermoFisher Scientific #A32723, 1:200 dilution) were diluted in the same blocking buffer, and sections were incubated with secondary antibody solution for 2 hours at room temperature. Sections were counterstained with Hoechst 33342 (ThermoFisher Scientific) for 1 hour at room temperature. PBS was used for all washes between staining steps. Coverslips were mounted with Prolong Gold antifade mounting media (ThermoFisher Scientific). Slides were imaged on an Olympus VS200 slide scanner, and images (.vsi format) were annotated in QuPath to identify cords with retraction of Sertoli cell apical processes.

### Testosterone Analysis

*Ex vivo* testicular testosterone production was assessed as described in our previous manuscript ^23^, using a method developed at the U.S. Environmental Protection Agency ^10^. Three testes were obtained from three different male fetuses in each litter and incubated in 500 µL of M199 media on an orbital shaker in a humidified cell culture incubator at 37°C and 5% CO_2_ for 3 hours. The media was then transferred to a -80°C freezer for storage until analysis. Media were analyzed by liquid chromatography-quadrupole time-of-flight-mass spectrometry (LC-QTOF-MS) using a procedure adapted from Yuan, Le, Wang and Li ^29^. In brief, 100 µL of each thawed media sample was first extracted by addition of 205 µL of acetonitrile containing testosterone-d3 internal standard and 1 mL of dichloromethane. The resultant organic layer was dried under nitrogen gas and then reconstituted with 100 µL of dichloromethane and derivatized with 10 µL of 10 mg/mL isonicotinoyl chloride hydrochloride in acetonitrile for 30 minutes. Following derivatization, samples were dried and reconstituted with 50 µL of methanol. For LC-QTOF-MS analysis, 10 µL of each reconstituted sample was injected.

LC was conducted using a 50 X 3mm Kinetex 2.6 µM C18 column on an ExionLC AC system. The mobile phase system consisted of water with 0.05% v/v formic acid and 50 mM ammonium formate (A) and 1:1 acetonitrile:methanol with 0.05% v/v formic acid (B). Separation of related sex hormones was achieved using a 14 minute gradient ranging from 2% to 98% mobile phase B, followed by a 2 minute re-equilibration. The flow rate was maintained at a constant 0.4 mL/min, and the column temperature was held at 30°C. QTOF-MS data collection was conducted using TOF-MS and TOF-MS/MS with information-dependent acquisition on a Sciex X500R QTOF-MS platform with ESI positive ionization. Quantification of media samples was performed by normalization of derivatized testosterone signal against testosterone-d3 internal standard followed by interpolation against an intra-batch linear calibration curve ranging from 50 to 3,200 ng/dL testosterone. Fragmentation spectra for isonicotinoyl chloride-derivatized testosterone, dehydroepiandrosterone, and testosterone-d3 are publicly available online at the MassBank of North America via accession numbers 0059502, 0059503, and 0059504 respectively.

### Phthalate Metabolite Analysis

Targeted analysis of dam serum was conducted by liquid chromatography-mass spectrometry. Serum samples stored at -80°C were allowed to thaw at ambient temperature prior to use. Samples were prepared by combining 100 µL of sample with 200 µL of acetonitrile, which were vortexed for one minute and incubated at 2-8°C overnight. Samples were then centrifuged using an Eppendorf 5424 benchtop centrifuge operated at 15,000 rpm for 15 minutes, and the supernatant was transferred to 300 µL amber glass autosampler vials for analysis.

The targeted analytical method was developed using standards for monobutyl phthalate (MBP), monoisobutyl phthalate (MIBP), monobenzyl phthalate (MBzP), monocyclohexyl phthalate (MCHP), mono-2-ethyl-5-hydroxyhexyl phthalate (MEHHP), mono-(2-ethylheyxl) phthalate (MEHP), monohexyl phthalate (MHP), and monopentyl phthalate (MPEP), as shown in Table 2. MBP and MIBP were not distinguishable from one another because they co-elute and maintain identical formula weights. Both analytes provided similar responses and ultimately MBP was chosen for calibration. Therefore, quantitative values reported for MBP may represent a mixture of MBP and MIBP. Analyses were performed using a ThermoFisher Scientific Q-Exactive HF-X Orbitrap mass spectrometer equipped with a heated electrospray interface and coupled to a Vanquish ultra-high performance liquid chromatography system. Separation of a 5 µL injection was performed using a ThermoFisher Hypersil Gold C18 column (2.1 x 100 mm, 1.9 µm). Mobile phases were prepared with 0.1% formic acid in LCMS-grade water (mobile phase A) and acetonitrile (mobile phase B). Column temperature was held at 60°C, and separation was performed at a constant flow rate of 0.4 mL/min with a gradient profile as follows: 30% B hold for 1 min, followed by a gradient ramp from 30% B to 95% B over 11 min and held for 2 min, and return to 30% B over 1 min and held for 2 min (total run time of 17 min). The mass spectrometer was operated in full scan mode while utilizing negative ionization with a resolution of 120,000 over a mass-to-charge range from 70 to 1050.

**Table 2.** Phthalate metabolites used for targeted serum analysis.

| phthalate metabolite | abbrev | CASRN | formula | parent C | total C | parent abbrev | source | PN | references |
| --- | --- | --- | --- | --- | --- | --- | --- | --- | --- |
| mono-isobutyl** | MIBP | 30833-53-5 | C<sub>12</sub>H<sub>14</sub>O<sub>4</sub> | 3 | 4 | DIBP | Accustandard | ALR-176S-CN |  |
| mono-n-butyl** | MBP | 131-70-4 | C<sub>12</sub>H<sub>14</sub>O<sub>4</sub> | 4 | 4 | DBP | Accustandard | ALR-135S-CN |  |
| mono-n-pentyl | MPEP | 24539-56-8 | C<sub>13</sub>H<sub>16</sub>O<sub>4</sub> | 5 | 5 | DPEP | Accustandard | ALR-177S-CN | Silva et al. <sup>81</sup> |
| monobenzyl | MBzP | 2528-16-7 | C<sub>15</sub>H<sub>12</sub>O<sub>4</sub> | 6 | 6 | BBP | Accustandard | ALR-134S-CN | Knox et al. <sup>82</sup> |
| monohexyl | MHP | 24539-57-9 | C<sub>14</sub>H<sub>18</sub>O<sub>4</sub> | 6 | 6 | DHP | Accustandard | ALR-175S-CN |  |
| monocyclohexyl | MCHP | 7517-36-4 | C<sub>14</sub>H<sub>16</sub>O<sub>4</sub> | 6 | 6 | DCHP | Accustandard | ALR-178S-CN | Silva et al. <sup>4</sup> |
| mono(2-ethyl-5-hydroxyhexyl) | MEHHP | 40321-99-1 | C<sub>16</sub>H<sub>22</sub>O<sub>5</sub> | 6 | 8 | DEHP | LGC Standards | TRC-M542510 | Koch et al. <sup>55</sup>, Knox et al. <sup>82</sup> |
| mono-(2-ethylhexyl) | MEHP | 4376-20-9 | C<sub>16</sub>H<sub>22</sub>O<sub>4</sub> | 6 | 8 | DEHP | Accustandard | ALR-138S-CN | Koch et al. <sup>55</sup> |
| monoheptyl (d4)* | MHEP | 24539-58-0 | C<sub>15</sub>H<sub>20</sub>O<sub>4</sub> | 7 | 7 | DHEP | LGC Standards | TRC-M546492 |  |

abbrev: abbreviation; parent C: length of parent side chain; total C: total C in side chain, including branches; PN: product number. \*Used as
internal standard. \*\*Identical formula weight.

### Proteomic Analysis

For proteomic analysis, 1-2 testes per litter (sample) were added to 200 µL T-PER Reagent with Halt Protease Inhibitor Cocktail, EDTA-Free (ThermoFisher), and homogenized using zirconium oxide beads in a Next Advance Bullet Blender on setting number 10 for 2 minutes. Total protein was quantified using the Pierce Rapid Gold BCA Protein Assay (ThermoFisher). 20 μg of protein was subjected to S-Trap Micro Digestion and Clean-up (Protifi) as specified by the manufacturer with modifications described here. Samples were lysed with 10% SDS, 50 mM triethylammonium bicarbonate buffer (TEAB) pH 7.5 and vortexed for 30 seconds. Proteins were reduced by incubation with dithiothreitol at 37°C for 45 minutes with mixing, alkylated by incubation with iodoacetamide at room temperature in the dark for 30 minutes, and then acidified by addition of phosphoric acid (2.5% final concentration) and vortexed for 30 seconds. 6 volumes of ice-cold binding buffer, consisting of 9 parts LC/MS grade methanol:1 part 1M TEAB, was added, and samples were transferred to S-Trap spin columns and centrifuged for 30 seconds at 4000 x *g*, then washed three times with binding buffer. Trypsin (Promega) in 50 mM TEAB was added to each micro spin column, and samples were incubated in a humidified chamber at 37°C overnight. Samples were cooled to room temperature for 15 minutes, 40 μL of 50 mM TEAB was added, and samples were centrifuged for 1 min at 4000 x *g*. Hydrophilic peptides were eluted with 40 μL of HPLC grade water and 0.1% formic acid (Fisher Scientific) and centrifuged for 1 min at 4000 x *g*. 40 μL of 50% LC/MS grade acetonitrile (Fisher Scientific) was added and allowed to stand for 5 minutes before centrifuging for 1 min at 4000 x *g*. Samples were concentrated using a chilled vacuum concentrator (Labconco), then reconstituted in 50 μL 5% acetonitrile and 95% water with 0.1% formic acid and spiked with indexed retention time peptides (Biognosys).

For data-dependent acquisition (DDA) LC-MS, approximately 200 ng reconstituted peptide samples were separated using a Vanquish Neo UHPLC on an Acclaim PepMap C18 Trap column and IonOpticks Aurora Ultimate Gen4 analytical column throughout a 60-minute gradient at 300 nL/min. Solvent A consisted of water with 0.1% formic acid, and solvent B consisted of 80% acetonitrile and 0.1% formic acid. The gradient consisted of loading/starting conditions of 5% B, 30% B over 50 min, 95% B over 5 min, then 99% B for 5 min. Combined control was used for loading and washing. Data was acquired on a Thermo Fisher Ascend Tribrid with spray voltage set to 2150 V, ion transfer tube temperature set to 275°C, and a user-defined lock mass of 445.12003. MS1 scans used wide quad isolation, orbitrap as the detector with a resolution of 120 K, mass range of 300-1500 m/z, maximum injection time of 50 ms, normalized AGC target of 250%, and RF Lens set to 30%. Data was collected in centroid mode with the following filters: MIPS (monoisotopic peak and peptide mode), charge state 2-7, precursor selection range 375-1500 m/z, dynamic exclusion (exclude after 1 time, 15 sec exclusion time, 10 ppm low and high mass tolerance, excluding isotopes), intensity range of 20,000 to 1E20 with relative intensity threshold of 20%. Data-dependent MS2 scans were acquired in centroid mode using cycle time 3 sec, normalized HCD fragmentation energy of 30%, and quadrupole isolation width of 0.7 m/z. Fragments were detected using the ion trap with turbo mode at a scan range of 150-2000 m/z, maximum injection time of 35 ms, normalized AGC target of 100%, and maximum injection time mode of dynamic was used.

For multiple accumulation precursor data-independent acquisition (MAP-DIA) proteomic analysis, approximately 200 ng of each reconstituted peptide sample was separated using a Vanquish Neo UHPLC on an Acclaim PepMap C18 trap column and IonOpticks Aurora Ultimate analytical column throughout a 60-min gradient at 300 nL/min. Solvent A consisted of water with 0.1% formic acid and solvent B consisted of 80% acetonitrile and 0.1% formic acid. The gradient consisted of loading/starting conditions of 5% B to 30% B over 50 min, 95% B over 5 min, and 99% B for 5 min. Data was acquired on a Thermo Fisher Ascend Tribrid mass spectrometer, using a method derived from Phlairaharn, Shannon, Zeng, Truong, Schoof, Ye and Searle ^30^. Briefly, the method consisted of a tSIM and tMSn scan. For the tSIM scans, quadrupole isolation and multiplex ions were selected with maximum number of multiplexed ions set to 6. Orbitrap resolution was set to 60K at a scan range of 350-1650 m/z. A maximum injection time of 10 ms and a normalized AGC target of 1000% was selected. RF lens was set to 40%, and the data type was set to centroid mode. For the tMSn scans, MSn level was set to 2, isolation utilized the quadrupole, with isolation widths of 16 m/z. HCD fragmentation was used with a normalized CE of 27%, and the detector was set to orbitrap at 30K resolution, RF lens at 40%. Maximum injection time was set to 60 ms, and the AGC target was 1000%. Data was collected in centroid mode.

.raw files were processed using Spectronaut (Version 21.0.260602.94842) with 1% peptide spectrum match, peptide, and protein FDR search constraints. All settings for spectral library generation using Pulsar were BGS defaults against the UNIPROT reference *Rattus norvegicus* protein .fasta (UP000002494_10116_20240602, 22,403 entries). Variable modifications consisted of oxidation (M), deamidation (N), Phosphorylation (STY), and protein N-terminal Acetylation. Carbamidomethylation was the sole static modification for the Trypsin search. PTM Localization was enabled using default probability thresholds (0.75), and single-hit proteins were excluded from analysis. The exported protein group report was further processed in Excel, and relative abundance was calculated as the intensity of each protein divided by the sum of all intensities for each given sample. The exported protein group report was further processed in Excel and Python 3.12 using an established pipeline ^31^, modified for use with Spectronaut reports, with the following dependencies: matplotlib (version 3.10.6), NumPy (version 2.3.3), SciPy (version 1.16.2), scikit-learn (version 1.7.2), Pandas (version 2.3.3), venny4py (version 1.0.3), and a custom Helpers package.

### Spatial RNA-seq

Spatial RNA-seq analysis of testes and mesonephroi that were snap-frozen and embedded in OCT was conducted using the Visium v1 Direct Placement Fresh Frozen assay (10x Genomics, Pleasanton, CA). All manufacturer protocols were followed. Briefly, blocks containing vehicle control (V) and high dose (A) phthalate mixture samples were prepared in OCT, with three biological replicate testes from different litters per block. This allowed for n=6 samples per group to be assayed using the four capture areas on the Visium slide. Blocks were trimmed on a Leica CM1960 cryostat to a depth that allowed all samples to be sectioned, and 10 µm sections were cut and placed on Visium tissue optimization or direct placement assay slides. For tissue optimization and gene expression assays, sections were methanol-fixed on the slide and stained with H&E according to Visium protocol CG000160. Tissue optimization was performed to identify optimal permeabilization time, according to Visium protocol CG000238. Prior to using blocks for the spatial assay, 10 x 10 µm sections from each block were used for RNA quality assessment. RNA samples were prepared from quality assessment samples using the PicoPure RNA Isolation Kit (ThermoFisher) and assessed on an Agilent 2100 Bioanalyzer (ThermoFisher). All quality assessment samples had RIN values of 8.3-9.8. Tissue sections placed on the Visium V1 assay slide were then used to prepare libraries, according to Visium protocols, using H&E staining. Briefly, sections were fixed with methanol, stained with H&E, and images were captured on an Olympus VS200 slide scanner. Sections were then permeabilized, RNA underwent first and second strand reverse transcription, and PCR amplification of cDNA. cDNA was cleaned up using SPRIselect beads (Beckman Coulter). cDNA was then fragmented, an adenine tail was appended to enable adaptor ligation, and samples were again cleaned up with SPRIselect beads. Final libraries with concentration ranging from 2.9 to 10.2 ng/µL were sent to the Molecular Biology Core Facility: Genomics at the Dana-Farber Cancer Institute for sequencing on an Illumina NovaSeq X sequencer in paired-end, 150 base pair format.

### Data Analysis and Statistics

Data derived from histological analysis, final quantities from serum metabolite analysis, and final testosterone values were summarized in Microsoft Excel, and statistical analysis was performed in Prism v10 or v11 (GraphPad Software, Boston, MA). To test for pairwise comparisons between the vehicle control and each dose level, either one-way ANOVA or the non-parametric Kruskall-Wallis test was used as follows. Residuals were tested for normal distribution using the D’Agostino-Pearson omnibus (K2) test. If residuals were not normally distributed, or if there was a greater than 4-fold difference between the smallest and largest dose-group variance within a dataset, the data were log transformed. If data were normally distributed and had equal variance, the were analyzed by one-way ANOVA, followed by Dunnett’s test to identify differences between any treatment group and the vehicle control. If log transformation did not resolve non-normality or unequal variance, data were analyzed using the nonparametric Kruskall-Wallis test, followed by Dunn’s multiple comparison test for each dose versus the vehicle control. Multiplicity-adjusted p-values (p-adj) were reported in the figures, along with individual data points and the mean ± SEM for each group. A comparison with p-adj < 0.05 was considered to be statistically significant. A dose-response analysis of serum metabolite concentrations was also performed in Prism, using the second-order polynomial (quadratic) regression equation.

Using the reported ED_50_ values for reduction of testosterone production by each of the phthalates in the mixture (Table 1), we predicted the dose-response for two apical endpoints using the dose addition equation reported by Rider and LeBlanc ^32^, where R is the response at a specific mixture concentration, based on the concentration, C_i_, for each chemical *i* with known EC_50_ and Hill slope, ρ.

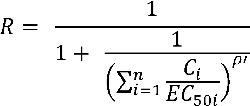

For comparison, we also calculated predicted dose-response for the two apical endpoints using the response-addition equation from Rider and LeBlanc ^32^.

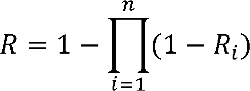

In the response-addition equation, the *R_i_* term is the predicted response for each individual chemical in the mixture.

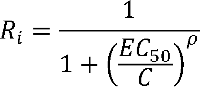

To calculate the dose-response for testosterone production, we set the maximum value to 100% of control. To calculate the predicted density of MNGs at each dose level, we used the inverse of the Hill slope from the testosterone production dose-response data and set the maximum value to 25.70 MNG/mm2, which was the highest group mean value from the current study, obtained in the B dose. To express the dose-response for the mixture in context of a well-characterized phthalate dose-response for induction of MNGs, we performed dose-response analysis using either the total phthalate dose (sum of all nominal phthalates in the dose mixture), or DPEP equivalent dose, based on relative potency factors (RPFs) calculated as the ratio of T production ED_50i_/T production ED_50DPEP_.

Benchmark Dose (BMD) analysis for phenotypic endpoints was performed using U.S. EPA Benchmark Dose Software, BMDS Online. Individual data for MNG density, *ex vivo* T production, seminiferous cord diameter, and proportion of cords with normal Sertoli cells (Sertoli cell score 0) were imported into BMDS. Best-fit model for each endpoint was determined using the maximum likelihood method, with the normal and non-constant variance assumptions. For all metrics, the benchmark response (BMR) was set to 10% change relative to control Models tested included all of the restricted exponential, Hill, polynomial, and power models, and the unrestricted linear model. Best-fit model was selected based on lowest Akaike information criterion (AIC). BMD_10_, BMDL_10_, and BMDU_10_ are reported based on the best fit model, using both DPEP equivalent and ∑phthalates (mg/kg/d) as the dose.

For analysis of proteomics data, the protein abundance matrix was subjected to quality control and filtering before performing differential expression analysis. For each pair-wise comparison, filtering was performed jointly across both conditions, i.e. vehicle control and dose groups, to exclude rows (proteins) in which fewer than 80% of samples had protein quantity values. The filtered data was subsequently used for differential protein abundance analysis using the R package limma ^33^. The resulting log2fold changes and adjusted p values (p-adj) were used to generate volcano plots using the ggplot function in R ^34^. Pairwise comparisons between vehicle control and any treatment group with p-adj < 0.05 were considered significantly differentially expressed (DE). All proteins that had at least one significant DE comparison were used for statistical overrepresentation analysis in PANTHER ^35^. The list of DE protein symbols from all treatment groups was uploaded as a single set, and overrepresentation was calculated, relative to the *R. norvegicus* genome reference list, using Fisher’s exact test with FDR adjustment for multiple testing. The mass spectrometry proteomics data have been deposited to the ProteomeXchange Consortium via the PRIDE ^36^ partner repository with the dataset identifier PXD085106.

Visium Spatial Gene Expression (v1) data were processed with Space Ranger v4.0.1 (10x Genomics) using spaceranger count with default settings. Reads were aligned to the mRatBN7-2-2024-A reference, and the tissue image for each capture area was used for automated fiducial alignment and tissue detection. Spatial barcodes and UMIs were used to quantify gene expression per spot. We delineated individual testes within each of four Visium capture areas, yielding 10 samples: 5 vehicle control testes and 5 testes treated with the phthalate mixture at dose A. For each spot, counts were scaled so that total counts summed to 10,000, and the resulting values were log1p transformed using Scanpy (v1.11.5). We used Conditional AutoRegressive-based Deconvolution (CARD) ^37^ for reference-based deconvolution to determine the cell type composition of each spot, with a published single-cell atlas of mouse fetal gonad development (E10.5–E16.5) (NCBI GEO accession no. GSE184708) ^38^ as the reference (18 annotated cell types). A mouse reference was used because no comparably annotated single-cell atlas of the fetal rat gonad was available. Deconvolution was performed on the 14,893 genes shared between the reference and the Visium data. Spatial niches were identified with CellCharter (v0.3.5) ^39^. Niches were defined using a Gaussian mixture model based on per-spot latent embeddings of spot expression aggregated across each spot’s neighborhood. We selected a 7-niche solution based on cluster stability and interpretability and named the niches following biological interpretation and cell-type enrichment. For each cell type, the CARD-estimated proportions were normalized within each niche so that all cell type proportions summed to 1 in that niche. Cell types with a maximum normalized proportion below 0.02 across all niches were excluded from the enrichment analysis, retaining 10 of the cell types identified by CARD. We computed cell type enrichment for each niche as the log2 ratio of the mean normalized proportion of a cell type within that niche to its mean proportion across the remaining niches. We derived p-values based on a one-sided Wilcoxon rank-sum test. Treatment effects on niche abundance were estimated by ordinary least squares (OLS) regression. To adjust for fraction of MT genes per spot and read depth, these variables were treated as covariates in the OLS regression. Differential expression between treatment groups was performed using the Wilcoxon rank-sum test on log-normalized counts across the full gene set, with Benjamini-Hochberg correction for multiple testing. We ran tests within defined sets of niches: N5 and N6 combined, N5 alone, N6 alone, and the Sertoli-enriched niches (N1, N2, N4). For gene-level comparisons summarized as per-sample values, we averaged spot-level expression within each sample. For poorly expressed genes, violin plots were restricted to spots with detected gene expression. Raw and normalized datasets are available from the NCBI SRA (PRJNA1531608) and GEO (GSE348356) databases.

## Results

### Dam and fetus weights and litter characteristics

Following *in utero* exposure to the 9-phthalate mixture, there were no dose-related differences in dam body weight on GD 21 (Fig. 1A) or change in dam body weight during the period of administration, GD 17-21 (Fig. 1B). The mean litter size was 13.4 fetuses across all 38 litters, and there was no significant difference in litter size by treatment group (Fig. 1C). The average M:F offspring sex ratio was 1.07 across all 38 litters, with no differences by treatment group (Fig. 1D). There were no groupwise differences in fetal body weight on GD 21 when analyzing all weights combined (Fig. 1E), male fetuses (Fig. 1F), or female fetuses (Fig. 1G). Individual data for these and other endpoints is reported in Table S1.

**Figure 1.**
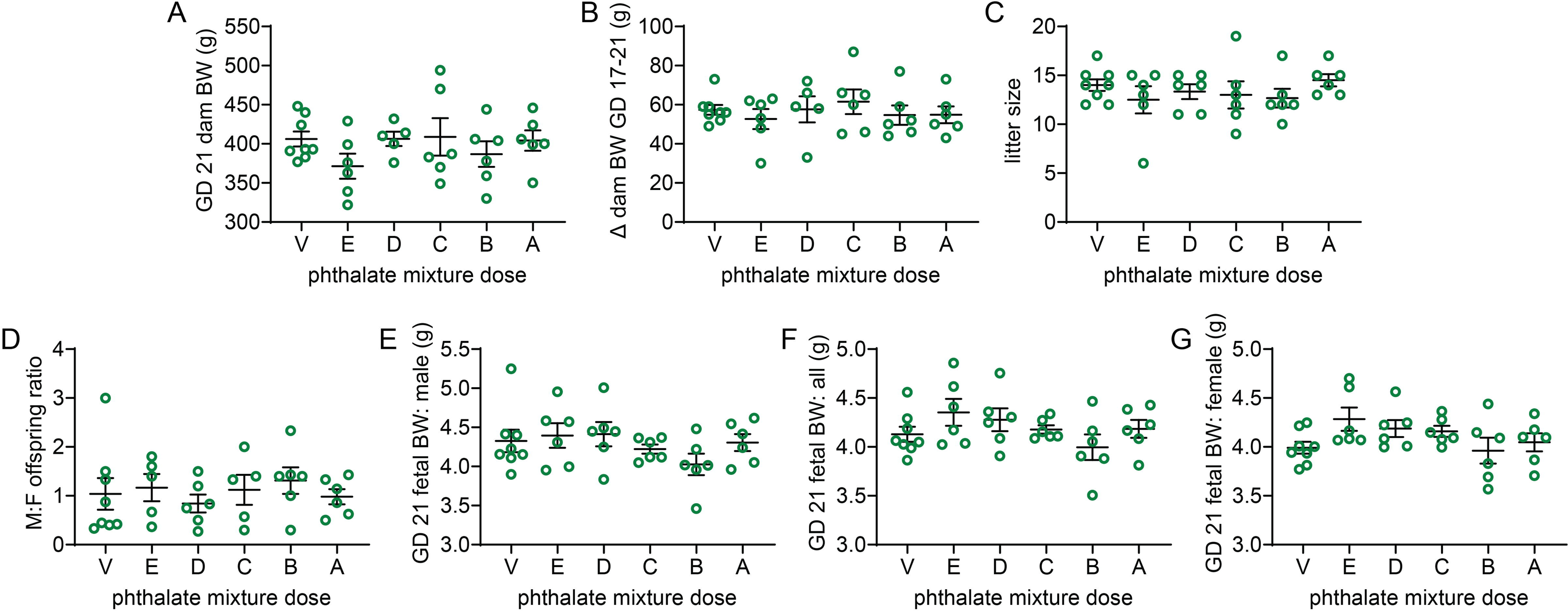
Dam and fetus weights and litter characteristics. **A**. Dam weight at time of euthanasia on GD 21. **B**. Change in dam weight from initiation of phthalate exposure on GD 17 to GD 21. **C**. Number of fetuses per litter. **D**. Male:female offspring sex ratio for all litters. **E-G**. Fetal body weights on GD 21 analyzed as all fetuses, regardless of sex (E), male fetuses (**F**), and female fetuses (**G**). There were no significant differences between any group and vehicle control (p > 0.05 for all statistical comparisons). Statistical comparisons were made by one-way ANOVA followed by Dunnett’s multiple comparison test in all cases except sex ratio, which was analyzed by Kruskall-Wallis test followed by Dunn’s multiple comparison test. Individual values are shown, overlaid with mean ± SEM.

### Maternal serum phthalate metabolites

All targeted phthalate metabolites that were detected in dam serum, and serum concentrations increased with increasing dose (Fig. 2). ∑ MBP + MIBP, MPEP, MHP, MEHP, MEHHP, and MBzP were detected in serum at all dose levels, while MCHP alone was not detected in dose E serum. Two vehicle samples had low but detectable levels of MHP, and one had a low but detectable level of MEHHP. Vehicle samples had no other detectable phthalate metabolites. Phthalate metabolite concentrations in dam serum increased monotonically across dose groups, both when they were analyzed individually and as the sum of all quantified metabolites. Serum concentrations of the sum of all metabolites were significantly greater than the vehicle control in doses C, B, and A (Kruskall-Wallis test with Dunn’s *post hoc* test). Quadratic regression of serum phthalate metabolite concentrations against total phthalate dose gave R^2^ values of 0.8945, 0.8848, 0.9368, 0.7071, 0.7387, 0.8653, 0.8380, and 0.8950 for ∑ all phthalate metabolites, ∑ MBP + MIBP, MBzP, MCHP, MEHHP, MEHP, MHP, and MPEP, respectively. However, notably, the data were non-normal for all metabolites except for MBP + MIBP, MHP, and MPEP. The order of metabolite concentration at a given dose level for the metabolites that were quantitatively measured was: [MBP + MIBP] > [MEHP + MEHHP] > [MPEP] > [MBzP] ≈ [MHP] > [MCHP] (Fig. 2), considering MBP and MIBP together because they are analytically inseparable, and considering MEHP and MEHHP together because they are both DEHP metabolites. This is roughly consistent with the order of parent compound concentration in the dosing mixture, which was DBP + DIBP + BBP (MBP + MIBP) > DHEP (not quantified) > DEHP (MEHP + MEHHP) > DIHEP (not quantified) > BBP alone (MBzP) > DPEP (MPEP) > DCHP (MCHP) > DHP (MHP) (Table 1).

**Figure 2.**
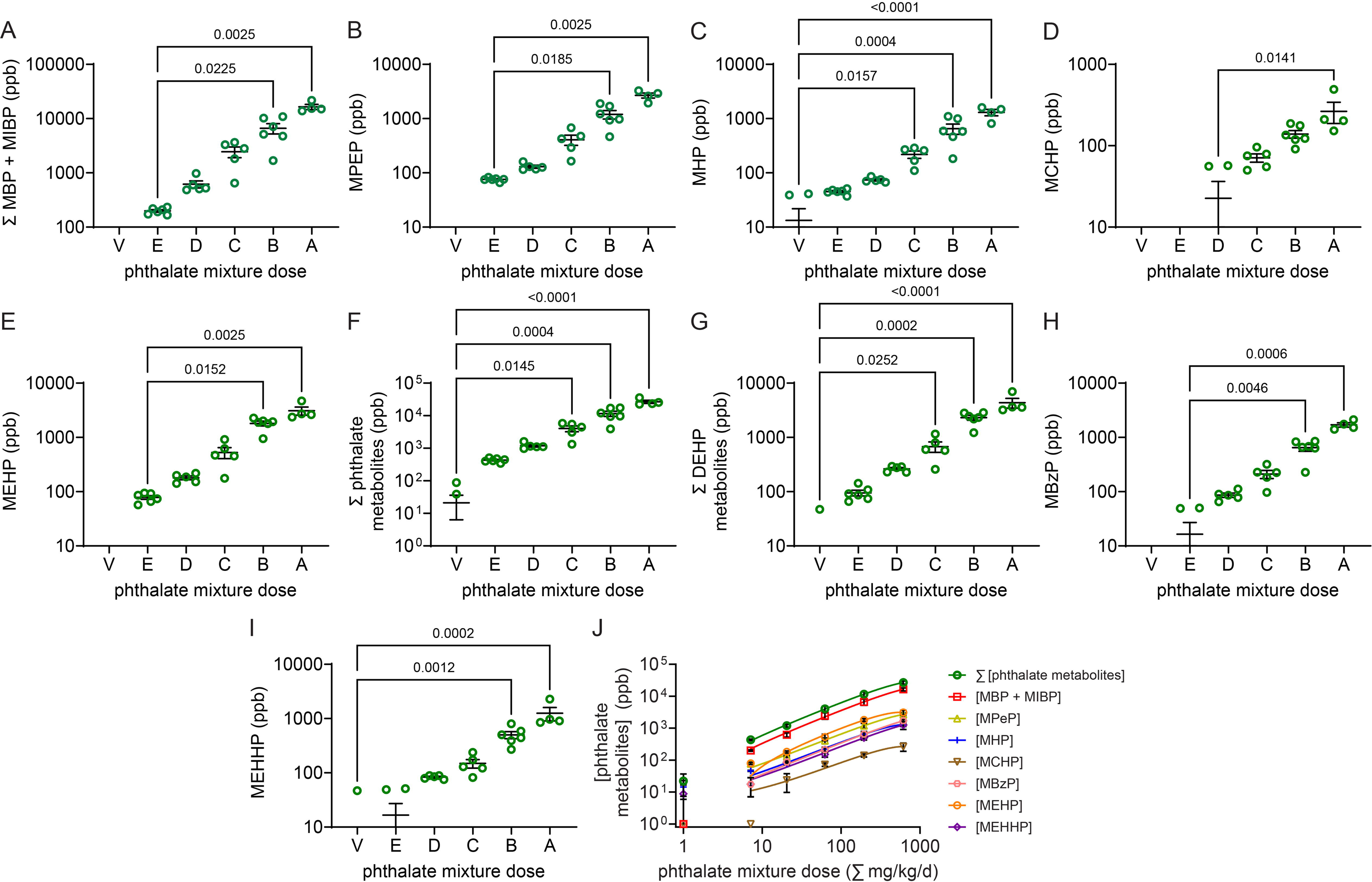
Analysis of dam serum. Targeted analysis was performed for 8 known phthalate metabolites by GC-MS/MS. Because MBP and MiBP are indistinguishable by formula weight, they are presented as a single value. Measured metabolites include **A**. ∑ MBP + MIBP, metabolites of DBP, DIBP, and BBP; **B.** MPEP, a metabolite of DPEP; **C.** MHP, a metabolite of DHP; **D.** MCHP, a metabolite of DCHP; **E.** MEHP and **F.** MEHHP, metabolites of DEHP; **G.** ∑ DEHP metabolites; **H.** MBzP, a metabolite of BBP. **I.** The sum of all phthalate metabolites from the targeted analysis. In **A-I**, statistical comparisons were made by Kruskall-Wallis test, followed by Dunn’s multiple comparison test. All metabolites were significantly greater in the than vehicle control in B and A dose animals, except for MCHP, which was significantly only in A. MHP was significant in C, and the sum of all metabolites was significant in C. Charts show adjusted p for each significant comparison. Individual values are plotted, overlaid with mean ± SEM. **J.** Quadratic regression analysis of all phthalate metabolites in dam serum. Metabolite concentrations increased monotonically across dose groups, with R^2^ values of 0.8945, 0.8848, 0.9368, 0.7071, 0.7387, 0.8653, 0.8380, and 0.8950 for ∑ all phthalate metabolites, ∑ MBP + MIBP, MBzP, MCHP, MEHHP, MEHP, MHP, and MPEP, respectively.

In addition to quantifying primary metabolites utilizing a targeted method, mass-to-charge ratios of nine suspected secondary metabolites, for which no reference standards were available, were extracted and peak area counts compared (Table S2). These secondary metabolites, including the DBP/DiBP metabolites MHBP/MHiBP, which have the same formula mass; the DPeP metabolites MHHP, MOPP, MCBP, MCPP, MCEP; the DEHP metabolites MECPP and MEOHP (probable); and the DIHEP metabolites MCHxP and MOHpP. Although we could not quantify these metabolites, they followed a similar pattern to the metabolites quantified in the targeted analysis, with the counts increasing on average with average dose. The only metabolites detected in the E dose group in the non-targeted analysis were MCHxP and MHBP/MHiBP. MECPP, MCHxP, and MEOHP were detected in the D dose group, while all of the non-targeted metabolites were detected in doses C, B, and A. This is consistent with these metabolites generally being present in lower quantities than the parent metabolites that were detected in the targeted analysis, possibly because they are secondary metabolites, produced later, and possibly more easily excreted. However, a detailed toxicokinetic analysis of this complex phthalate mixture has not been performed. There was no other known metabolite of dicyclohexyl phthalate that could explain why MCHP was not detected in the E dose in the targeted analysis, which may suggest that DCHP is more slowly metabolized by the esterase than the phthalates with linear or branched aliphatic side chains.

### Multinucleated germ cells and seminiferous cord morphology

Testes exposed to the phthalate mixture differed from vehicle control samples in seminiferous cord structure, including the morphology of germ cells and Sertoli cells. The density of MNGs in H&E-stained testis sections was elevated following exposure to the 9-phthalate mixture (Fig. 3A). MNG density was significantly greater in doses B and A than vehicle control (p-adj < 0.0001 and p-adj = 0.0004, respectively, Kruskall-Wallis test with Dunn’s *post hoc* test), with the greatest control mean of 25.70 MNGs/mm^2^ in dose B, and a similar mean of 19.47 MNGs/mm^2^ in dose A (Fig. 3B). The dose C mean of 5.842 MNGs/mm^2^ was not significantly greater than the vehicle control (p-adj = 0.1084). The mean MNG density in doses E and D was very low (0.1793 and 0.5660 MNG/mm^2^, respectively) and not statistically significant. The average number of nuclei/MNG did not differ significantly by group, but was highest in dose A, with a mean of 2.727 nuclei/MNG (Fig. 3C). To test whether this apparent increase in nuclei per MNG resulted in an increase in the total density of nuclei in MNGs, here referred to as MNN, we analyzed MNN by Kruskall-Wallis test, followed by Dunn’s test. MNN were significantly greater than vehicle control in dose B (p-adj = 0.0003) and A (p-adj = 0.0022) but not dose C (p-adj = 0.3046) (Fig. 3D). For comparison with measured dose-response, we modeled the predicted mixture dose response using the dose-addition and response-addition equations (Fig. 3E). The dose (independent variable) used for dose-response modeling was DPEP equivalent dose, which was derived using relative potency factors (RPFs) for each chemical in the mixture (Table 3). The predicted ED_50_ for MNG induction was 35.66 mg/kg/d DPEP equivalent by the dose-addition equation and 84.17 mg/kg/d DPEP equivalent by the response-addition equation, compared to the measured ED_50_ of 23.64 mg/kg/d DPEP equivalent, and the DPEP ED_50_ of 27.70 mg/kg/d that we reported previously ^23^.

**Figure 3.**
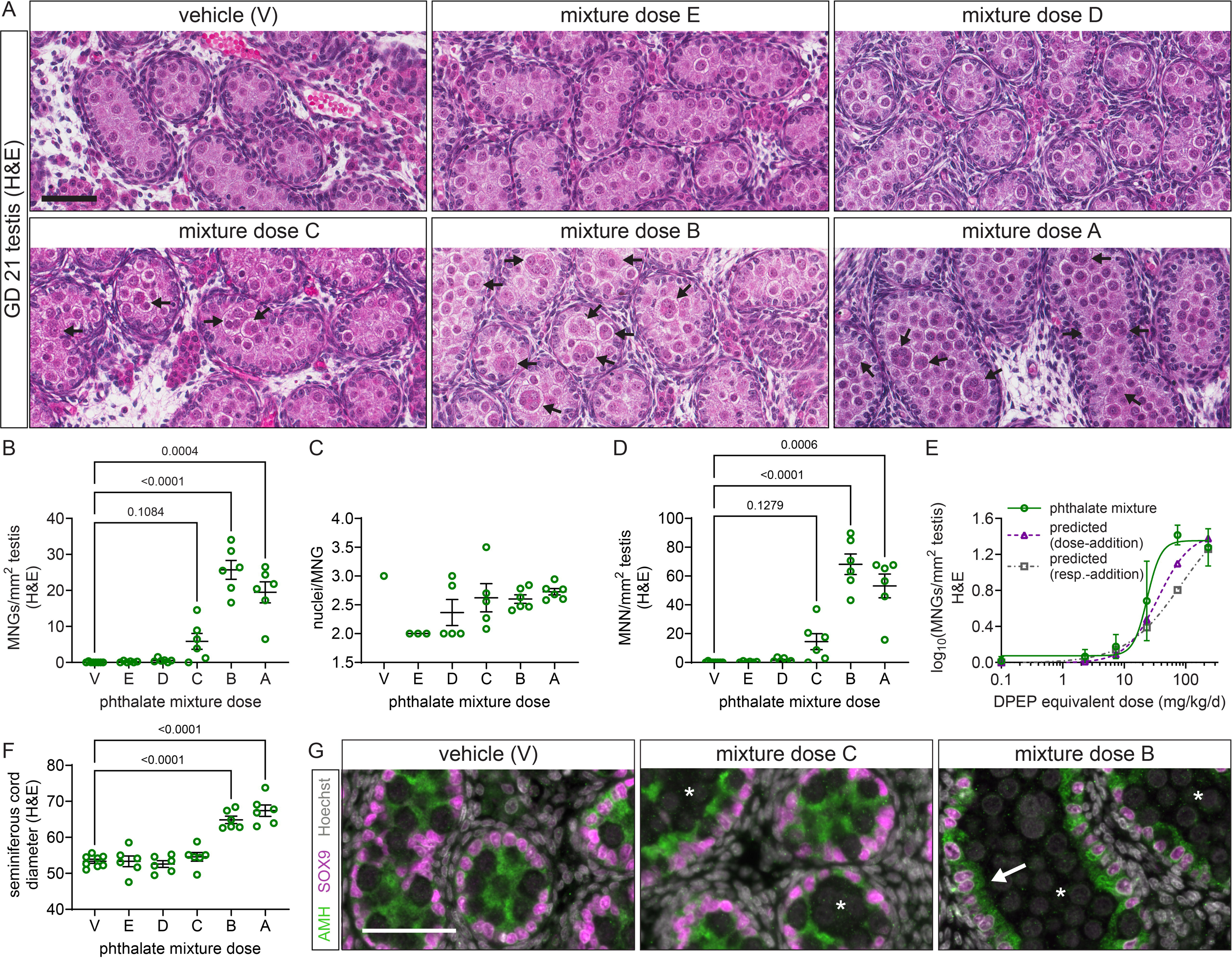
Multinucleated germ cells (MNGs) and seminiferous cord morphology. **A.** Representative images of hematoxylin and eosin (H&E)-stained paraffin sections of gestational day (GD) 21 fetal testes following exposure to the 9-phthalate mixture or vehicle control. Arrows: MNGs. Scale bar = 60 μm. **B.** MNG density in histological sections (Kruskall-Wallis test followed by Dunn’s test). **C.** Number of nuclei per MNG (Kruskall-Wallis test followed by Dunn’s test). **D.** Density of nuclei in MNGs (MNN) per mm^2^ cross-sectional area. Charts in **B-D** show individual replicate values with mean ± SEM. (n = 6-8 litters/group). Brackets are labeled with adjusted p-value. **E.** Dose-response analysis of log_10_MNGs/mm2 on H&E sections was performed in Prism using nonlinear regression, with DPeP equivalent dose (mg/kg/d) as the independent variable. Vehicle control value was set to 0.1 mg/kg/d to enable log transformation of dose on the x-axis, and one pseudocount was added to all values to allow for log transformation of 0 values. Model constraints: ED_50_ > 0. Purple dashed line shows the dose-response predicted using the dose-addition equation. Gray dashed line shows the dose-response predicted using the response-addition equation. Predicted ED_50_ was 35.66 mg/kg/d by dose addition and 84.17 mg/kg/d by response-addition; measured ED_50_ was 23.64 mg/kg/d. **F.** Mean GD 21 seminiferous cord diameter following exposure to the 9-phthalate mixture or vehicle control, quantified on H&E sections (one-way ANOVA followed by Dunnet’s test). **G.** Representative images of paraffin sections of gestational day (GD) 21 fetal testes following exposure to the 9-phthalate mixture or vehicle control, immunostained for SOX9 (Sertoli cell nuclei) and AMH (Sertoli cell cytoplasm), with Hoechst counterstain. Vehicle-treated sample shows normal cord morphology with Sertoli cell processes in the center of the cord, surrounding germ cells. Phthalate mixture -treated samples show regions of seminiferous cords with moderate (dose C) or severe (dose B) retraction of Sertoli cell processes (asterisks). Phthalate mixture dose B-treated sample shows a seminiferous cord region with a flattened, cuboidal-like Sertoli cell layer (arrow). Scale bar = 50 μm.

**Table 3.** Dose-addition modeling of MNGs and ex vivo T production.

| outcome | non-linear regression<br>DPeP eq. (mg/kg/d) |  |  |  | non-linear regression<br>Σphthalates (mg/kg/d) |  |  |  |
| --- | --- | --- | --- | --- | --- | --- | --- | --- |
|  | ED<sub>50</sub> | R<sup>2</sup> | predicted ED<sub>50</sub><br>(dose-addition) | predicted ED<sub>50</sub><br>(response-addition) | ED<sub>50</sub> | R<sup>2</sup> | predicted ED<sub>50</sub><br>(dose-addition) | predicted ED<sub>50</sub><br>(response-addition) |
| MNGs<br>(H&E) | 23.64<br>(17.63, 29.31) | 0.888 | 35.66 | 84.17 | 62.59<br>(46.70, 77.59) | 0.888 | 94.33 | 220.2 |
| ex vivo T<br>(LC-QTOF-MS) | 40.55<br>(12.47, 147.2) | 0.484 | 81.72 | 157.7 | 107.2<br>(33.6, 389.0) | 0.484 | 216.0 | 417.3 |

eq: equation; LC-QTOF-MS: liquid chromatography-quadrupole time-of-flight-mass spectrometry; MNGs: multinucleated germ cells; T: testosterone.

In addition to induction of MNGs, seminiferous cord diameter, quantified on H&E-stained sections, increased with increasing phthalate dose (Fig. 3F). Samples treated with the B or A doses had significantly greater seminiferous cord diameter on average. Mean diameter in the V, E, D, and C dose groups was between 52.59 and 54.58 µm. In the B and A doses, mean seminiferous cord diameter was 64.85 and 67.42 µm, respectively, similar to what we have reported previously for DBP ^17^ and DPeP ^23^. Changes in seminiferous cord diameter were accompanied by altered localization of AMH in Sertoli cells, suggestive of changes in Sertoli cell cytoplasmic structure (Fig. 3G). Using immunofluorescent labeling of SOX9 to identify Sertoli cell nuclei and AMH to label Sertoli cell cytoplasm, we observed that seminiferous cords with normal Sertoli cell morphology were present in all samples and predominated in vehicle-treated samples. However, in samples treated with dose C and above, we also observed seminiferous cords with moderately retracted Sertoli cell apical projections, resulting in cords that had no AMH signal in the center of the cord, as well as cords with severely retracted projections in which the Sertoli cells had a flat, cuboidal epithelium-like apical surface.

### Reduction of fetal testicular testosterone production

*Ex vivo* T production was significantly lower than vehicle control level in testes exposed to the B and A doses of the phthalate mixture (p-adj = 0.0207 and 0.0018, respectively, one-way ANOVA with Dunnett’s *post hoc* test) (Fig. 4). The control mean value of 674.6 ng/dL was reduced to 180.0 ng/dL in dose A, a reduction of 73.3%. The ED_50_ was 40.55 mg/kg/d DPEP equivalent, compared to 81.72 mg/kg/d DPEP equivalent predicted by the dose-addition equation, 157.7 mg/kg/d DPEP equivalent predicted by the response-addition equation, and 134.1 mg/kg/d DPEP measured in our prior experiment ^23^.

**Figure 4.**
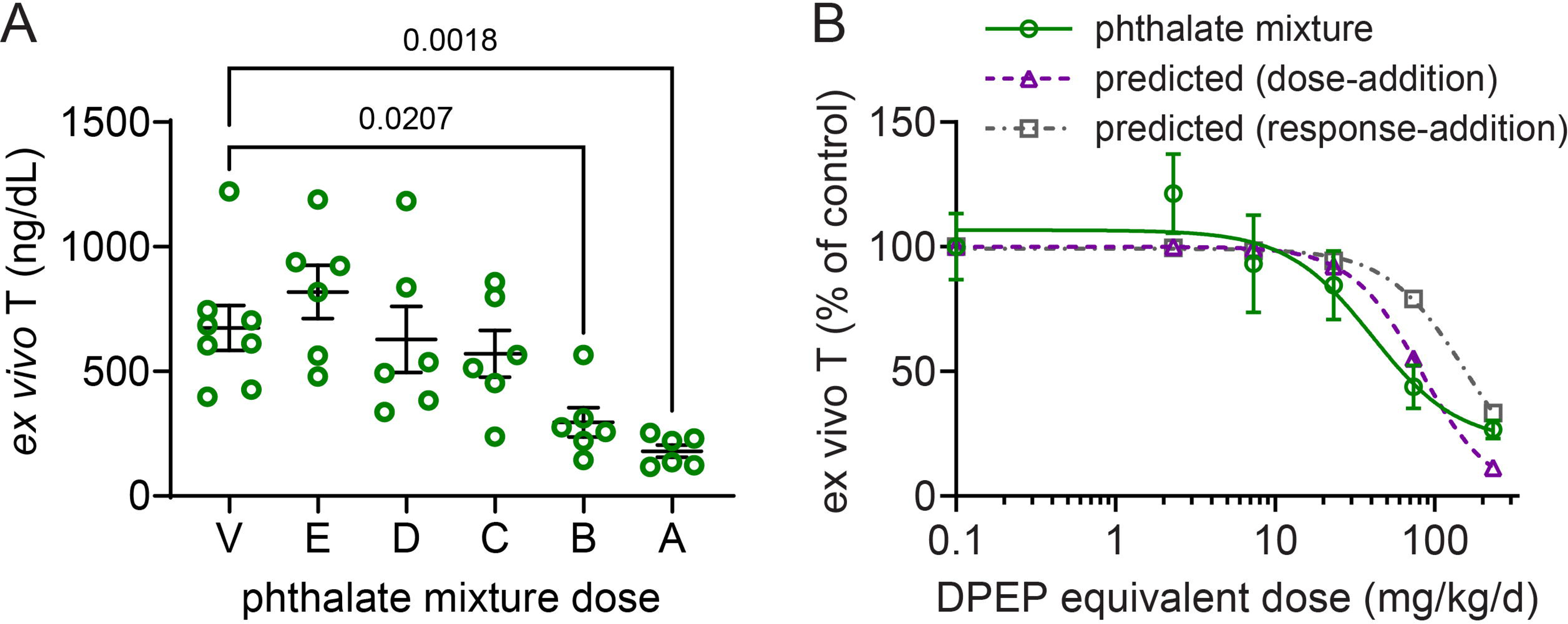
Ex vivo testosterone production in GD 21 rat fetal testes. **A.** *Ex vivo* testosterone (T) production (one-way ANOVA followed by Dunnett’s test). Chart shows individual replicate values with mean ± SEM (n = 6-8 litters/group). Brackets are labeled with adjusted p-value. **B.** Dose-response analysis of *ex vivo* T production data was performed in Prism using nonlinear regression (inhibition, four-parameter) with DPeP equivalent dose (mg/kg/d) as the independent variable. Data normalized to % of concurrent control mean. Vehicle control value was set to 0.1 mg/kg/d to enable log transformation of dose on the x-axis. Model constraints: ED_50_ > 0, bottom > 0. Purple dashed line shows the dose-response predicted using the dose-addition equation. Gray dashed line shows the dose-response predicted using the response-addition equation. Predicted ED_50_ was 81.72 mg/kg/d by dose-addition and 157.7 mg/kg/d by response-addition; measured ED_50_ was 40.55 mg/kg/d.

### Benchmark dose modeling of apical endpoints

MNG density, *ex vivo* T production, seminiferous cord diameter, and Sertoli cell morphology data were used for BMD modeling in EPA BMDS software (Table 4). The lowest BMD for any endpoint was MNG density, with a BMD_10_ (BMDL, BMDU) of 3.517 (2.275, 4.919) mg/kg/d DPEP equivalent or 9.402 (6.093, 13.13) mg/kg/d total nominal phthalate dose. The BMD_10_ values in DPEP equivalent dose were 13.826, 28.922, and 39.097 for *ex vivo* T production, Sertoli cell morphology, and seminiferous cord diameter, respectively. The 95% confidence intervals overlapped for all three of endpoints, but none overlapped with the BMD_10_ for MNG density.

**Table 4.** Benchmark dose modeling results.

| outcome | BMDS analysis:<br>DPeP eq. (mg/kg/d) | | BMDS analysis:<br>Σphthalates (mg/kg/d) | | dose at BMD<sub>10</sub> (mg/kg/d) | | | | | | | | |
| --- | --- | --- | --- | --- | --- | --- | --- | --- | --- | --- | --- | --- | --- |
|  | BMD<sub>10</sub><br>(BMDL, BMDU) | best fit<br>model | BMD<sub>10</sub><br>(BMDL, BMDU) | best fit<br>model | DIBP | DBP | BBP | DPEP | DCHP | DHP | DIHEP | DEHP | DHEP |
| MNGs<br>(H&E) | 3.517<br>(2.275, 4.919) | exp 5* | 9.402<br>(6.093, 13.13) | exp 5* | 1.39 | 0.76 | 0.83 | 0.39 | 0.30 | 0.29 | 1.75 | 1.74 | 1.94 |
| SC diameter<br>(H&E) | 39.097<br>(32.995, 53.072) | Hill | 103.469<br>(87.321, 135.52) | Hill | 15.34 | 8.40 | 9.18 | 4.34 | 3.28 | 3.15 | 19.25 | 19.15 | 21.38 |
| ex vivo T<br>(LC-QTOF-MS) | 13.826<br>(5.214, 28.436) | Hill | 36.639<br>(15.5, 78.798) | Hill | 5.43 | 2.98 | 3.25 | 1.54 | 1.16 | 1.12 | 6.81 | 6.78 | 7.57 |

Benchmark dose (BMD) modeling conducted in EPA Benchmark Dose Software (BMDS), based on both DPEP equivalent dose (DPeP eq.) and total phthalate dose. BMD<sub>10</sub>: BMD corresponding to a benchmark response (BMR) of 10% change from control, assuming normality and non-constant variance. Dose at BMD<sub>10</sub> is the proportional dose of each phthalate in the exposure mixture at the model-derived BMD<sub>10</sub>. Models were selected based on lowest Akaike information criterion (AIC). \*questionable model fits: models with control stdev fit > 1.5 and nonconstant variance test failed.

### Phthalate mixture effects on fetal testis proteome

Following exposure to the 9-phthalate mixture, changes in fetal testis protein expression were detected in samples from the A, B, and C doses, versus vehicle control, using proteomic profiling by mass spectrometry (Table S3). Each group was compared to the vehicle control using Limma, after filtering out rows with more than one missing value. Of 7306 proteins that were detected in any sample, 5790, 5227, 5967, 5224, and 5466 remained after filtering in groups A, B, C, D, and E, respectively. The total number of differentially expressed proteins (p-adj < 0.05), relative to the vehicle control, was 34 (2 upregulated, 32 downregulated), 33 (5 upregulated, 28 downregulated), 4 (0 upregulated, 4 downregulated), 0, and 0, in groups A, B, C, D, and E, respectively (Fig. 5, Table S4). There was strong dose-dependency, with 4 proteins significant at all three dose levels and 19 significant in the A and B doses, while 11 proteins were significant only in dose A and 10 only in dose B. Many of the DE proteins have known expression localization. Of 44 total proteins, at least 31 are known to be expressed in Leydig cells, 16 in Sertoli cells, and 14 in germ cells; at the subcellular level, 12 DE proteins are known to be localized in the endoplasmic reticulum and 11 in the mitochondrion (Table S4). We performed a functional enrichment analysis in PANTHER to determine the Gene Ontology (GO) Biological Processes (BPs) and Cellular Components (CCs) that were enriched in the DE protein list (Table 5, Table S5). The enriched CCs included the mitochondrion and the endoplasmic reticulum. Enriched BPs were almost exclusively related to cholesterol, steroid, and steroid hormone biosynthesis and metabolism, but also included carboxylic acid biosynthetic process, cellular detoxification, modified amino acid metabolic process, fatty acid metabolic process, and alcohol catabolic process, among others.

**Figure 5.**
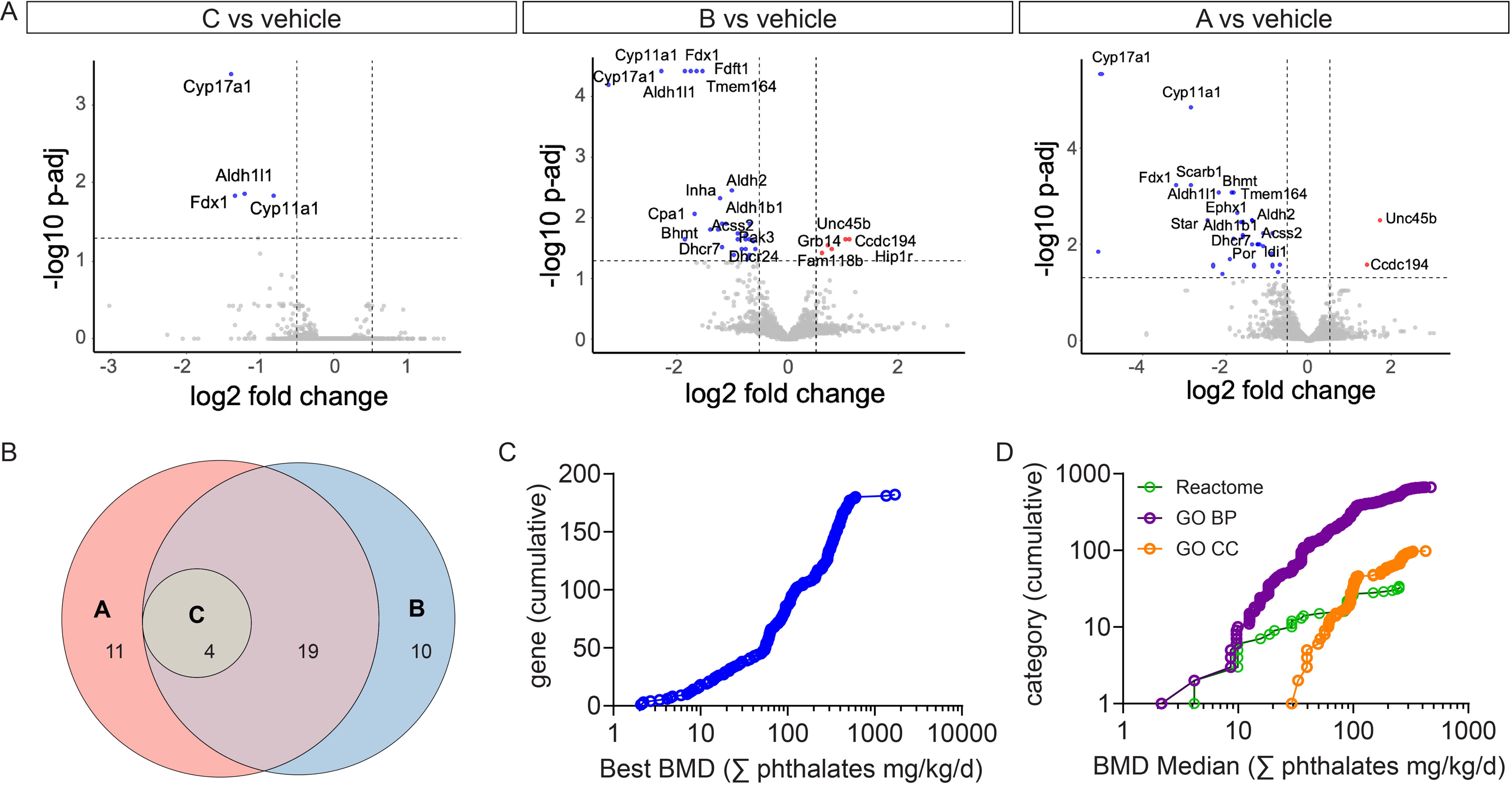
Effect of 9-phthalate mixture on GD 21 fetal testis proteome. **A**. Limma analysis of proteomic data identified 4, 33, and 44 differentially expressed (DE) proteins in GD 21 fetal testes exposed to the C, B, and A doses of the 9-phthalate mixture, respectively (p-adj < 0.05). No DE proteins were detected in comparisons of D or E dose vs. vehicle. Blue dots: downregulated proteins; red dots: upregulated proteins. **B.** Venn diagram of DE proteins in each treatment group vs. vehicle control comparison. **C.** Cumulative density plot showing the best BMD for the 182 proteins that passed the Williams’ trend test filter. **D.** Cumulative density plot showing BMDmedian for each significantly enriched Gene Ontology (GO) biological process (purple), GO cellular component (orange) and Reactome pathway (green).

**Table 5.**
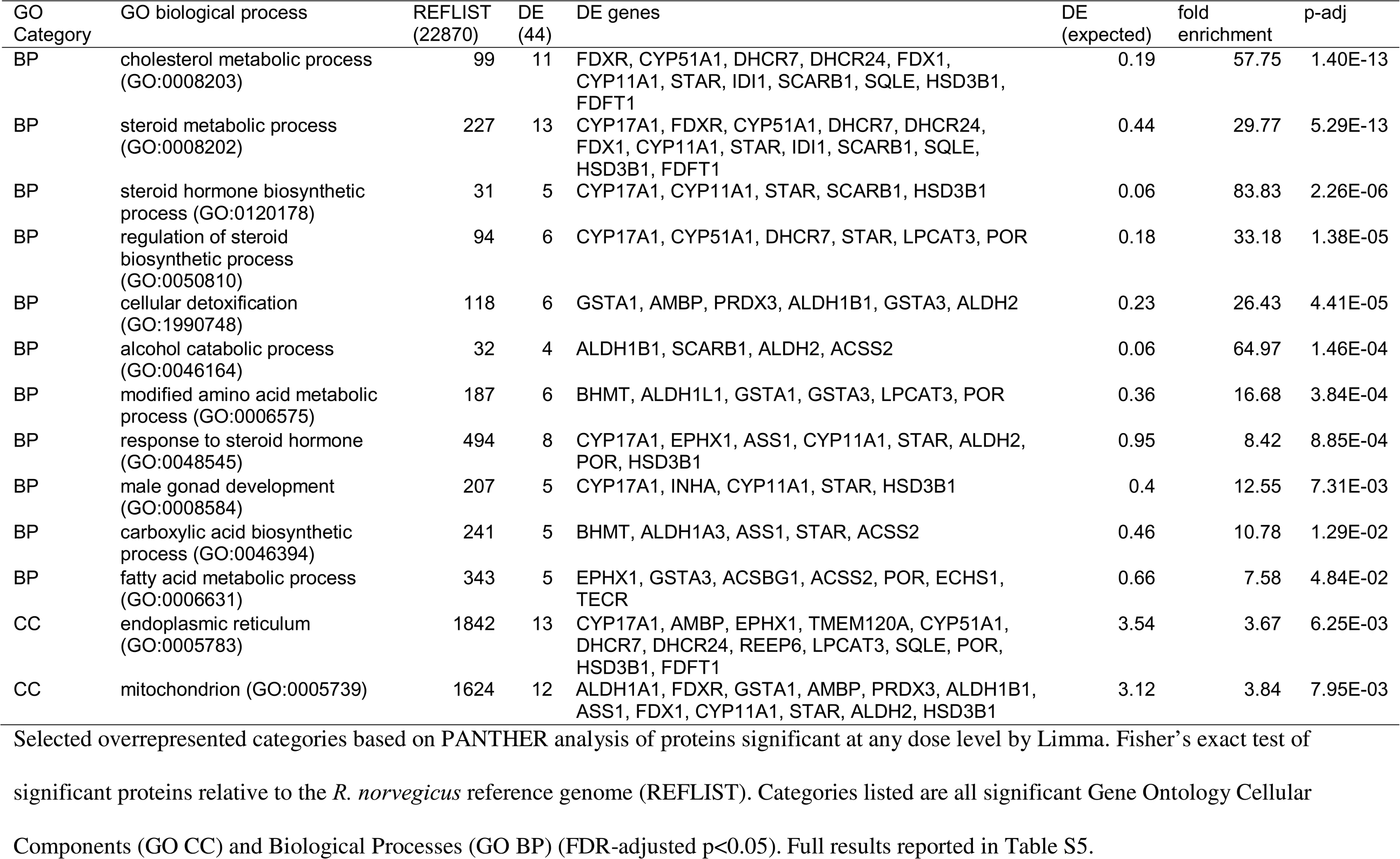
Functional enrichment analysis of DE proteins.

Proteomic BMD analysis was conducted in BMDExpress3. After removing one sample from group A with many missing values and filtering to remove any rows with missing values, 3369 rows were analyzed in BMDExpress3. A less stringent protein-level approach was used in BMD analysis than in Limma, and 182 proteins passed the Williams trend test prefilter (unadjusted p<0.05, fold change ≥ 1.1) (Table S6. The 182 proteins that passed the prefilter were used for BMD Analysis (Table S7). The most common best model fit was linear (37.4%), followed by polynomial 2 (29.1%), Hill (24.7%), exponential 5 (7.7%), and power (1.1%). The lowest protein-level BMD in the dataset was for SCARB1 at 2.07 (0.204, 14.074) mg/kg/d total phthalate, by the exponential 5 model. Protein-level BMD values for all 182 proteins were used for category analysis to identify BMDmedian values for overrepresented GO BPs and CCs and Reactome pathways (Table 6, Table S8). There were 668 GO BPs, 98 GO CCs, and 34 Reactome pathways with at least 3 genes that passed all filters. There was a clear trend toward overrepresentation of categories related to cholesterol and steroid hormone biosynthesis among the categories with the lowest BMDmedian values. However, the overall lowest BMDmedian was 2.16 mg/kg/d total phthalates for GO:0051701, biological process involved in interaction with host. The lowest steroid-related term was GO:0042446, hormone biosynthetic process, with a BMDmedian value of 4.15 mg/kg/d, based on the downregulation of FDX1, STAR, and SCARB1.

**Table 6.**
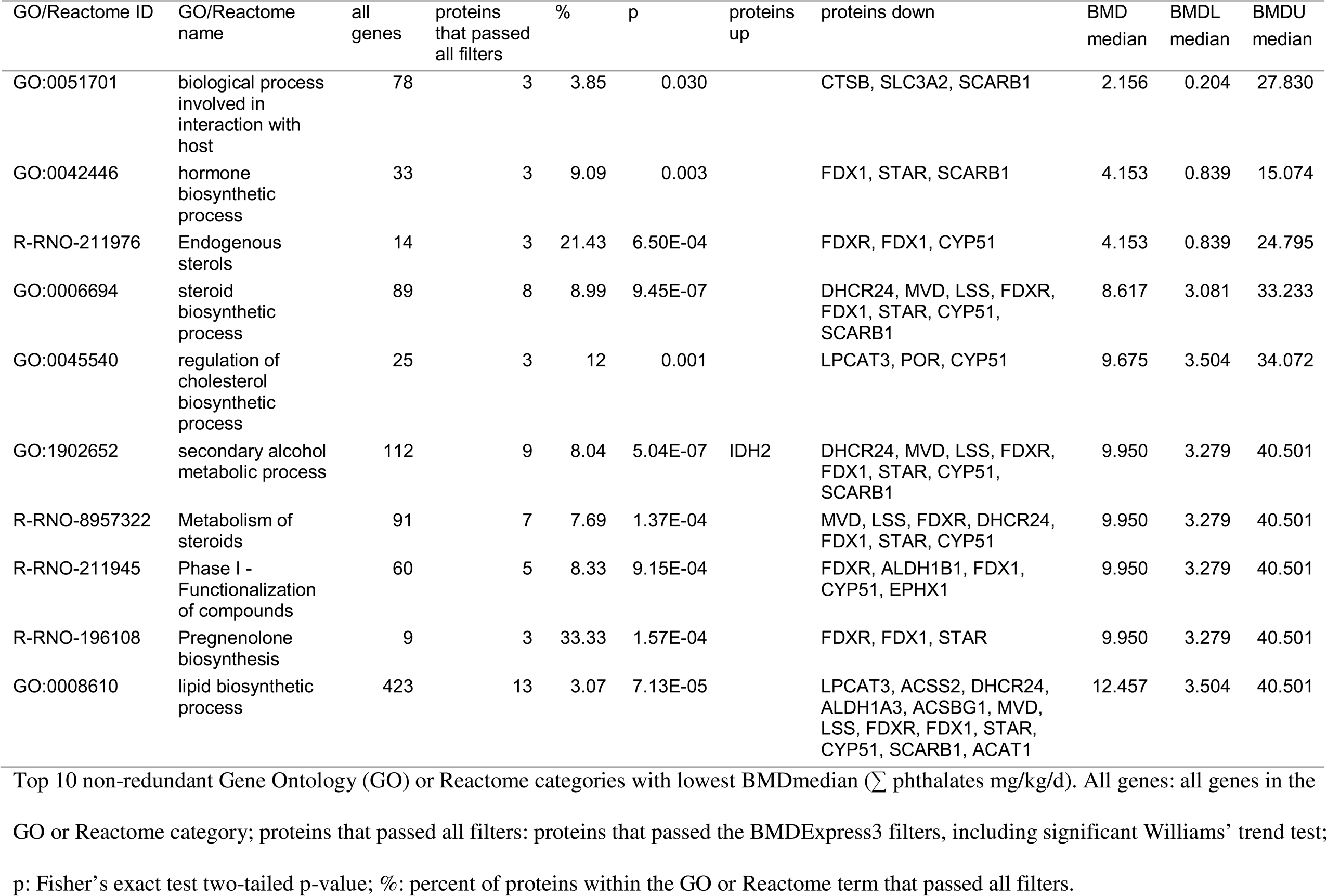
Proteomic Benchmark Dose Analysis.

### Spatial gene expression effects of phthalate mixture

We performed spatial RNA-seq analysis of GD 21 testes following treatment with vehicle or the phthalate mixture dose A from GD 17-21 (n=5 per group). In spatial RNA-seq analysis, we identified seven niches (Fig. 6A-B) based on the spatial neighborhood composition of cell types estimated by reference-based deconvolution (CARD) (Table S9). Niches are composed of multiple spots, in turn composed of multiple cells each due to the 55 µm spot size and 10 µm section thickness. Based on the relative presence of mesonephric cells, endothelial cells, perivascular cells, blood cells, germ cells, Sertoli cells, interstitial progenitors, immune cells, peritubular myoid cells (PTMCs), and fetal Leydig cells, spots were classified as N0, mesonephric/PTMC-enriched; N1, seminiferous cord; N2, cord and associated vasculature; N3, peritubular myoid cell/tunica; N4, vascular/mixed interstitium; N5, fetal Leydig cell; and N6, interstitial progenitor. Proportionally, phthalate treatment led to a decrease in the number of spots in N5 and N6, the two Leydig cell-containing niches, and an increase in all other niches (Fig. 6C, Table S10). In all niches, including niches composed solely of Leydig cells, *Cyp17a1*, *Star*, *Scarb1*, *Hsd3b1*, *Cyp11a1*, and *Fdx1* were downregulated, consistent with phthalate-driven suppression of steroidogenic and cholesterol biosynthetic activity in Leydig cells (Fig. 6D-F). The expression of Leydig cell genes was largely unchanged in N5 alone, but significantly decreased in N6, with the exception of *Hsd3b1* being apparently upregulated in N5 (Fig. 6G). This suggests that following the loss of steroidogenic gene expression under phthalate exposure, some Leydig cells may have essentially been reclassified as interstitial precursors, rather than as Leydig cells. In the Sertoli cell-containing niches, N1, N2, and N4, two of the most significant changes were downregulation of *Vim* and upregulation of *Amh* (Fig. 6H, Table S10), consistent with an effect of phthalate exposure on differentiation of fetal Sertoli cells.

**Figure 6.**
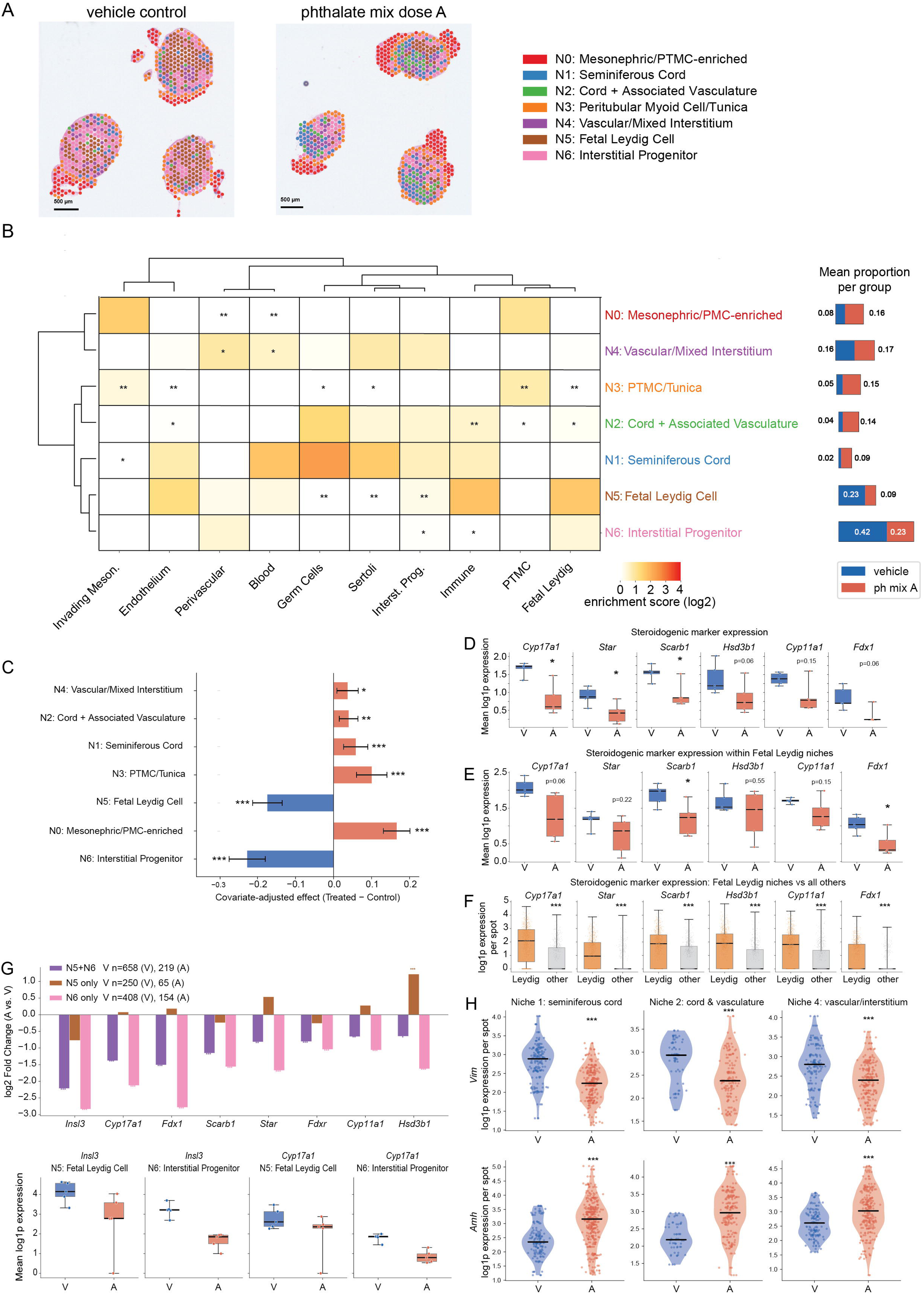
Effect of 9-phthalate mixture on spatial gene expression in GD 21 fetal testis. **A.** Niche overlays for a subset of control and phthalate mixture dose A-treated GD 21 rat fetal testes. Each spot was assigned a niche N0-N6 based on the expression of cell type-specific markers. **B.** Heatmap showing the enrichment of each fetal testicular cell type in each niche and the proportion of spots corresponding to each niche in vehicle and A dose groups. *p<0.05, **P<0.01, ***p<0.001 by one-sided Wilcoxon rank-sum test (niche vs. all other niches). **C.** Chart showing the relative change in the proportion of spots corresponding to each niche in phthalate mixture dose A vs. vehicle. *p<0.05, **p<0.01, ***p<0.001 by ordinary least-squares regression. **D.** Expression of Leydig cell marker genes involved in steroidogenesis and cholesterol biosynthesis in all niches. *p<0.05 by two-sided Wilcoxon rank-sum test on per-sample mean expression. **E.** Expression of Leydig cell marker genes in the fetal Leydig cell-containing niches only. *p<0.05 by two-sided Wilcoxon rank-sum test on per-sample mean expression. **F.** Relative expression of Leydig cell marker genes in Leydig cell niches (N5 and N6) vs. all other niches. ***p<0.001 by one-sided Wilcoxon rank-sum test. **G.** Relative expression of Leydig cell marker genes in control vs. phthalate mixture dose A-treated testes, broken down into each Leydig cell-containing niche, N5, N6, and N5+N6 combined. *p-adj<0.05, **p-adj<0.001, ***p-adj<0.001 by Wilcoxon rank-sum test on log-normalized counts across the full gene set, with Benjamini-Hochberg correction for multiple testing. **H.** Violin plots showing expression of *Vim* and *Amh* in vehicle and phthalate mixture dose A-treated samples within the Sertoli cell-containing niches, N1, N2, and N4. ***p<0.001 by Wilcoxon rank-sum test on non-zero spot-level expression values.

## Discussion

We previously reported that induction of multinucleated germ cells (MNGs) is a sensitive, quantitative metric of the toxic effects of phthalates on the fetal seminiferous cords in general and the germ cells in particular ^23^. In this experiment, we found that the toxicity of a phthalate mixture to testis morphogenesis was dose-additive, using MNGs as the quantitative adverse outcomes to test this hypothesis and *ex vivo* T production as a more data-rich comparator (Fig. 3 and 5, Table 3). In both cases, the experimental data were sufficient to support dose-response and BMD modeling, with the dose range covering two doses at or near maximum effect and two statistical no-effect doses. Using the dose addition model, we predicted an ED_50_ for T production of 81.72 mg/kg/d DPEP equivalent or 216.0 mg/kg/d ∑phthalates. The measured ED_50_ for T production was lower than the predicted value, but the confidence interval overlapped the predictions in both cases, at 40.55 (12.47, 147.2) mg/kg/d DPEP equivalent or 107.2 (33.16, 389.0) mg/kg/d ∑ phthalates. For MNG density, the measured ED_50_ of 23.64 (17.63, 29.31) mg DPEP equivalent/kg/d or 62.59 (46.70, 77.69) mg/kg/d ∑ phthalates was very close to the dose-addition model prediction of 35.66 or 94.33 mg/kg/d, respectively. In both cases, the ED_50_ estimated by the dose-addition model was approximately 1.5- to 2-fold higher than the measured ED_50_, while the response-addition model predicted ED_50_ values approximately 3.5- to 4-fold higher than the measured values. The strong concordance between dose-addition prediction and observed data for MNGs is evidence that phthalate effects on fetal germ cells are dose-additive. Moreover, these results are congruent with prior reports that phthalates and other anti-androgens have dose-additive effects on testosterone and androgen-dependent endpoints ^24-26^. The measured T production values in the present study had higher variance than MNG density, which may have contributed to the greater difference between observed and predicted values for T production than for MNGs. Despite the fact that the MNG and T production dose-addition predictions were both derived from prior T production data, the dose-addition model-based ED_50_ prediction was lower for MNGs than T production. This resulted from the fact that T production data has a downward adverse direction and was bounded in our analysis such that the minimum value was > 0, a constraint that does not apply to the increasing MNG density values.

We also calculated benchmark dose (BMD) for apical endpoints using BMDS software, and we found that the lowest BMD_10_ was for MNG density (Table 4). This is consistent with the effects of DPEP in Gupta et al. ^23^; however, in the present experiment, the confidence intervals were non-overlapping, allowing us to conclude that MNG density was a more sensitive phthalate response within the parameters of the current study. As we noted previously, the most sensitive window for T production is during the masculinization programming window, which is earlier than the GD 17-21 exposure window used in the present study. Therefore, it is possible that the effect of the phthalate mixture on T production would be more potent if an earlier window of exposure was used. We used the late exposure window because it encompasses the quiescent period of germ cell development, during which the testis is sensitive to MNG induction ^17^. Seminiferous cord diameter is known to be greater on average in phthalate-treated samples than controls. In our previous experiment, this effect was not as sensitive a response to phthalates as either MNG density or T production, and there was no difference in the density of Sertoli cells per unit area ^23^. A phthalate-induced difference in Sertoli cell proliferation and density has been reported when the exposure window begins earlier in gestation ^16^. As in our previous experiment, in the present study there was a larger average seminiferous cord diameter in phthalate mixture-treated testes. Because we expected no difference in Sertoli cell density, we assessed Sertoli cell morphology to determine whether we could identify a phenotypic change in the Sertoli cells that precedes (or occurs at a lower dose than) the increased seminiferous cord diameter. Because of previous reports that phthalate treatment alters the morphology of Sertoli cell apical processes ^15^, we used a combination of AMH and SOX9 IF to identify cords where the Sertoli cell processes were retracted and therefore not visible in the center of the cord and between germ cells (Fig. 3G). We observed seminiferous cords with retracted apical Sertoli cell processes in dose C, and more severe retraction in doses B and A. These results reinforce that fetal Sertoli cells are sensitive to phthalate exposure, and that morphologic changes in Sertoli cells co-occur with phthalate-induced MNGs. We have now observed that MNGs are induced at lower doses of phthalates than the dose required to reduce T production in studies of a phthalate mixture (Fig. 4, Table 4) and of DPEP ^23^. Because phthalates are known to induce MNGs in mice and rats, as well as human fetal testis tissue ^20, 40^, this suggests that fetal Sertoli and germ cell toxicity of phthalates is relevant to human health.

In BMD analysis of the phthalate mixture proteomic data, using BMDExpress3 (Fig. 5, Table 6), the lowest BMDmedian value was GO: 0051701: biological process involved in interaction with host, at 2.156 mg/kg/d, and the lowest BMDmedian for a term with >7 proteins that passed all filters was GO:0006694: steroid biosynthetic process, at 8.617 mg/kg/d, measured as ∑all phthalates, placing the proteomic BMD close to the lowest apical BMD_10_ of 9.402 (6.093, 13.13) ∑all phthalates, for MNG density. A proteomic POD based on the lowest BMDmedian value for this dataset would be approximately 4.36-fold lower than the lowest apical BMD_10_, and therefore health-protective. This is consistent with past reports on the consistency of apical and transcriptomic points of departure ^41-45^, Although the majority of published work on molecular points of departure (mPODs) to date has utilized transcriptomic data, and sometimes metabolomic data ^46, 47^, the present analysis indicates that proteomic data could be similarly useful for this purpose, despite the fact that the depth of information is generally lower for proteomic analysis than for transcriptomic analysis.

To date, no molecular initiating event has been identified that can explain the toxicity of phthalates to the fetal testis, despite the fact that the metabolism of phthalates and the cellular changes and apical outcomes of phthalate toxicity are well-characterized for some phthalates ^7^. Parent phthalate diesters are cleaved by intestinal or hepatic esterases to produce monoesters that are toxicologically active ^48-50^. For high-production volume phthalates, such as DBP and DEHP, pregnant rat toxicokinetics have been described. Placental transfer of the primary monoester metabolite is efficient, such that fetal serum phthalate metabolite exposure can be as high as 80% of the maternal serum exposure ^49^. Many of the metabolites that we report here (Fig. 2) have been detected in human biomonitoring data, including MBP, MBzP, MEHP, MEHHP, MEOHP, MECPP, MiBP, and MCPP, which are metabolites of DBP, BBP, DEHP, DiBP, and DPP, respectively ^4, 6, 51-54^. MBP, MiBP, MEHHP, MEOHP, and MEHP have been confirmed as metabolites in human volunteers following ingestion of the respective parent compounds, DBP, DiBP, and DEHP ^55, 56^. Because of the elevated DEHP exposure associated with blood transfusions in the NICU, MEHP, MEOHP, MEHHP, and MECPP have all been detected in neonates following blood transfusions, with the order of both AUC in the first 60 hours and T_1/2elimination_ following transfusion being MECPP > MEHHP and MEOHP > MEHP ^5^. In this experiment, we detected all of the metabolites for which we had quantitative standards in approximately the same rank order of serum concentration as the rank order of parent compounds in the dosing mixture, with the exception that MBzP and MCHP appeared slightly lower in the rank of measured metabolites than the ranks of the respective parent compounds in the mixture. This may suggest that phthalates with ring structures in the side chain rather than simple or branched aliphatic side chains are metabolized more slowly. This should be addressed in future studies of phthalate mixture toxicokinetics. We also detected many metabolites for which we did not have standards for quantification (Table S2). Because of the rapid biotransformation of phthalates, it is not known whether the secondary metabolites exert toxicity or what is their potency relative to the primary monoester metabolite, but animal studies have clearly established that *in vivo* administration of the primary monoester metabolites results in the characteristic effects of the parent diesters ^57-62^. Notably, human biomonitoring studies have also confirmed that humans are exposed to metabolites of short-chain phthalates such as monoethyl phthalate and monomethyl phthalate ^4^, as well as long-chain phthalates such as diisononyl phthalate and diisodecyl phthalate ^63^. These parent compounds were excluded from the mixture in the present study because of evidence that they are not toxic to the rat fetal testis in the case of MEP and MBP ^10, 19^ and because of a relative paucity of dose-response data in the cases of DINP and DIDP. However, DINP and DIDP are emerging phthalates, and there is evidence that DINP exerts fetal testicular toxicity ^64^.

This analysis of the effects of phthalates on the fetal testis proteome and spatial transcriptome provide new insights about phthalates’ toxic mode of action in the fetal testis. We chose these two analyses to address unexplored aspects of the phthalate response. Although there are a significant number of prior publications describing bulk transcriptomic effects of phthalates, there is relatively little data on phthalate effects at the proteome level or on the spatial or cell type-specific effects of phthalates. In a previous analysis, we found that the bulk transcriptomic effects of DPEP were largely dose-dependent and were not confined to Leydig cell-expressed genes, with more DE genes known to be expressed in the Sertoli cell than any other cell type ^23^. However, functional enrichment analysis coalesced around the known effects of phthalates on the steroid hormone biosynthetic pathway. Here, the proteome data show some of the same patterns and some differences (Tables 5-6). In the phthalate mixture-treated proteome, there was an overwhelming enrichment of downregulated proteins that are expressed in the mitochondria and/or endoplasmic reticulum and that have roles in cholesterol or steroid metabolism (Table S5). There was also enrichment of numerous fatty acid or lipid transport and metabolism terms and the icosanoid metabolic process term, as well as the Leydig cell differentiation and male gonad development terms. The male gonad development term and many of the steroid and cholesterol-related terms consist of downregulated proteins including INSL3, STAR, CYP17A1, CYP11A1, HSD3B1, SCARB1, and SQLE, which are a core group of known phthalate-responsive genes from transcriptomic analyses ^65-68^. This adds to evidence that cholesterol metabolism and transport, which are upstream of steroid hormone metabolism, are core targets of phthalates ^23^. There is evidence from other experimental systems that phthalates alter energy metabolism. For example, phthalates cause toxicity in the liver ^69^ by acting as agonists of peroxisome proliferator activated receptors (PPARs) ^70, 71^. Activation of PPARs is considered to be the mechanism by which phthalates induce cancerous and pre-cancerous hepatic and pancreatic lesions, as well as interstitial adenomas in the testis, in the rat and mouse ^72, 73^. Accordingly, phthalates alter fatty acid metabolism ^74^, and this is reflected in enrichment of fatty acid metabolism as a category in the proteomic data, including downregulation of EPHX1, GSTA3, ACSBG1, ACSS2, POR, ECHS1, and TECR (Table 5). Consistent with the ability of phthalates to act as PPARA ligands, in one experiment, a mixture of phthalate metabolites increased ATP production in cultured granulosa cells at 24 and 72 h. However, there is evidence that PPAR activation or signaling is not involved or is not the only mechanism of testicular toxicity of phthalates in the fetal testis ^75-77^.

In spatial RNA-seq analysis comparing vehicle control and phthalate mixture dose A (Fig. 6), the Leydig cell signature was relatively high in two niches, which we named the Fetal Leydig cell (N5) and Interstitial Progenitor (N6) niches. One of the biggest changes detected in the analysis was a decrease in the proportion of spots assigned to N5 and N6 in phthalate-treated samples (Fig. 6C), while the within-cluster differential expression data showed downregulation of Leydig cell-expressed steroidogenic genes across all niches (Fig. 6D) and within the fetal Leydig cell niches in particular (Fig. 6E). This suggests that the phthalate treatment suppressed expression of Leydig cell gene expression, as expected, and may have even caused certain spots containing Leydig cells to be classified as non-Leydig cell niches, given that the steroidogenic genes are markers of Leydig cell identity. This likely resulted in an apparent decrease in Leydig cell-containing spots, despite prior IF evidence from DPEP-treated testes that the density of HSD3B1-positive Leydig cells was not changed by phthalate treatment in late gestation ^23^.

In relation to sex hormone signaling, it is especially notable that INHA was significantly downregulated following phthalate exposure. INHA forms heterodimers with INHB subunits to form inhibin A and B. In a recent experiment, mice with a deletion for the *Inha* gene, which results in low inhibin expression and high activin A, had reduced testicular androstenedione and T, as well as germ cell phenotypes including reduced fetal germ cell density, ectopic germ cells, and MNGs ^78^, very similar to the fetal testicular phenotype that follows from phthalate exposure in the fetal rat testis. In the present study, the proteomic analysis identified reduced INHA levels in the A and B doses (Table S4), consistent with a previous study showing downregulation of *Inha* in the rat fetal testis interstitium after DBP exposure ^79^. This pattern and the correlation with the *Inha* knockout mouse study suggests that low *Inha* expression and elevated activin A levels contribute to phthalate toxicity.

The remaining, non-Leydig cell niches in the spatial RNA-seq analysis were proportionally overrepresented in the phthalate-treated samples (Fig. 6A). There were two Sertoli cell-containing niches that were of interest. In the Sertoli cell containing-niches, which we called Cord and Associated Vasculature (N2) and Seminiferous Cord (N1), as well as a mixed cell type niche that we called Vascular/Mixed Interstitium (N4), both *Amh* and *Vim* were DE (Fig. 6H). *Vim* was down-regulated following phthalate exposure, consistent with prior published evidence that DBP caused retraction of the vimentin filaments in rat fetal Sertoli cells ^15^. *Amh* was upregulated in the Sertoli cell-containing niches. In the proteomics analysis, AMH expression was not significantly changed, while AMH localization patterns by IF (Fig. 3G) differed between phthalate-treated samples and vehicle controls in that apical Sertoli cell processes were retracted in phthalate-treated samples. It is possible that *Amh* mRNA expression is elevated in the Sertoli cells of phthalate-treated rats because of a delayed Sertoli cell maturation phenotype or that the reduced *Inha* expression in phthalate-treated fetal Sertoli cells could lead to elevated activin A and therefore elevated FSH, which could alter *Amh* expression ^80^. Based on the available data, we can only hypothesize as to the mechanisms by which phthalates alter *Amh* expression, but the increase in *Amh* mRNA, coupled with the altered Sertoli cell phenotype, is more evidence that phthalates alter fetal Sertoli cell maturation and function.

In summary, we report for the first time that a phthalate mixture-induced germ cell toxicity in the rat fetal testis was both dose-dependent and dose-additive. Consistent with our prior analysis of DPEP toxicity ^23^, the quantitative effect of phthalates with the lowest ED_50_ and BMD_10_ was induction of MNGs. Analysis of proteomic and spatial transcriptomic data confirmed prior knowledge of phthalate toxicity, including enrichment of differentially expressed (downregulated) proteins involved in steroid hormone biosynthesis. However, we also report that there was broad disruption of fatty acid, lipid, icosanoid, and amino acid metabolism, and enrichment of proteins expressed in the mitochondria and endoplasmic reticulum, which presents several avenues for future research to determine the upstream processes responsible for initiation of phthalate toxicity. Benchmark dose analysis based on proteomic data produced similar BMD estimates to those derived from apical effects, indicating that proteomic BMDs may be useful for derivation of PODs.

## Acknowledgments

The authors thank Justin M. Conley, PhD, for technical assistance with study design and implementation of the dose-addition model. The authors thank Nicholas DaSilva, PhD, for proteomic analysis and David Silverberg for embedding and cutting of histological specimens.

